# Polygenic risk score for Alzheimer’s disease and trajectories of cardiometabolic risk factors in children

**DOI:** 10.1101/580548

**Authors:** Roxanna S Korologou-Linden, Linda O’Keeffe, Laura D Howe, George Davey Smith, Hannah Jones, Emma L Anderson, Evie Stergiakouli

**Affiliations:** Medical Research Council Integrative Epidemiology Unit (IEU) at the University of Bristol, University of Bristol, Bristol, UK; Population Health Sciences, Bristol Medical School, University of Bristol, Bristol, UK; Centre for Academic Mental Health, Population Health Sciences, Bristol Medical School, University of Bristol, Bristol, UK; School of Oral and Dental Sciences, University of Bristol, Bristol, UK

**Keywords:** Polygenic risk scores, Alzheimer’s disease, modifiable risk factors, cardiometabolic, metabolic, inflammatory, ALSPAC

## Abstract

**INTRODUCTION:** Cardiometabolic factors are implicated in the aetiology of Alzheimer’s disease and may lie on the pathways linking genetic variants to Alzheimer’s disease across the life course. We examined whether polygenic risk scores (PRS) were associated with cardiometabolic health indicators through childhood and adolescence.

**METHODS:** In 7,977 participants from the Avon Longitudinal Study of Parents and Children, we tested whether a PRS for Alzheimer’s disease was associated with trajectories of cardiometabolic risk factors. We examined trajectories for height 1-18 years; lean and fat mass 9-18 years; systolic and diastolic blood pressure 7-18 years; glucose and C-reactive protein 9-18 years; insulin 10-18 years; high and low-density lipoproteins and triglycerides birth-18 years. We also examined birthweight, interleukin-6 (IL-6) at age 9 years and physical activity at ages 11, 12, and 15 years.

**RESULTS:** No consistent associations were observed between the PRS excluding genetic variants in the apolipoprotein E (*ApoE*) gene region and cardiometabolic factors trajectories across childhood and adolescence.

**CONCLUSION:** We did not detect evidence to suggest that the PRS for Alzheimer’s disease acts through childhood and adolescent cardiometabolic risk factors. Further studies should examine whether these associations emerge later in adulthood when variation in cardiometabolic risk factors is likely to be greater.

## INTRODUCTION

There is growing evidence from observational studies for a role of midlife hypertension, obesity, hyperlipidaemia [1], hyperglycaemia [2], and inflammation [3, 4] in the aetiology of Alzheimer’s disease. Evidence for a role of cardiometabolic risk factors in later life is less clear as observed associations are often either null or protective [1, 5, 6]. This age-dependent change in the direction of the associations observed for cardiometabolic risk factors has been suggested to potentially reflect selection bias, as fatal cardiometabolic disease events would reduce the risk of being diagnosed with Alzheimer’s disease, which usually occurs late in life [7]. Another possible explanation for these findings is reverse causation, due to the potential presence of undiagnosed Alzheimer’s disease (which has been shown to be associated with a lowering of blood pressure [5] and weight loss [6]) at the time of assessment of the cardiometabolic risk factors.

Large genetic consortia have identified common genetic variants associated with late-onset Alzheimer’s disease [8], with the ε4 allele of the *ApoE* gene conferring the greatest risk, increasing the risk up to twelvefold [9]. *ApoE* adversely affects lipid profiles [10] and elevates the risk of coronary artery disease [11]. Although the genetic variants (apart from *ApoE4*) individually increase risk by a small amount, they can be summarised to construct a polygenic risk score (PRS) for Alzheimer’s disease which has a good ability to correctly classify those with and without the disease (area under the receiver operating characteristic curve [AUROC] = 78.2% with a P-value threshold of ≤0.5) [12]. Apart from *ApoE*, the relationship of these genetic variants to cardiovascular disease is uncertain.

PRS can be used to investigate potential prodromal phenotypes and or pathways through which the variants influence Alzheimer’s disease risk. There are very few studies [13, 14] examining the association between cardiometabolic risk factors in early life, likely due to the long follow-up required across the life course. We tested the hypothesis that a PRS for Alzheimer’s disease (beyond the *ApoE* gene) acts through cardiometabolic pathways in children and adolescents.

## METHODS

### Participants

The Avon Longitudinal Study of Parents and Children (ALSPAC) is a prospective birth cohort which recruited pregnant women with expected delivery dates between April 1991 and December 1992 from Bristol UK [15, 16]. 14,541 pregnancies were initially enrolled, from which 13,867 live births occurred in 13,761 women. Including two later rounds of recruitment there were 15,445 eligible children. Information on health and development of children and their parents were collected from regular clinic visits and completion of questionnaires. Research clinics were held when the participants were approximately seven, nine, 10, 11, 13, 15, and 18 years old. The study website contains details of all the data that is available through a fully searchable data dictionary: http://www.bris.ac.uk/alspac/researchers/data-access/data-dictionary/. Ethical approval was obtained from the ALSPAC Law and Ethics Committee and the Local Ethics Committees.

### Cardiometabolic measures

#### Anthropometry

Birthweight was obtained from obstetric records. At the clinics between 4 months and 5 years, crown-heel length was measured using a Harpenden Neonatometer and from 25 months onwards, standing height was measured using a Leicester Height Measure. From age 7 years, all children were invited to yearly clinics, where measurements of standing height were obtained using the Harpenden Stadiometer. At each clinic at ages nine, 11, 13, 15, and 18 years, whole body less head, and central fat and lean mass were derived from whole body dual energy X-ray absorptiometry (DXA) scans evaluated using a Lunar prodigy narrow fan beam densitometer.

#### Cardiometabolic and inflammatory factors

At each clinic (ages seven, nine, 10, 11, 12, 15, and 18), systolic and diastolic blood pressure were measured at least twice. The mean of two measurements was used in the analysis. Insulin was obtained from cord blood at birth. Non-fasting glucose was also measured at age seven as part of the metabolomics assays, using Nuclear Magnetic Resonance Spectrometry (NMR). Fasting glucose and insulin were also available for a random 10% of the cohort at age nine years, as well as research clinics at 15 and 18 years. Triglycerides, high-density lipoprotein (HDL-c), and total cholesterol (TC) were available from cord blood and venous blood, subsequently. Non-fasting samples were available from research clinics at ages seven and nine years; fasting measures were available from clinics at 15 and 18 years. Non-HDL-c was calculated by deducting HDL-c from total cholesterol. Hence, the trajectories of triglycerides, HDL-c, non-HDL-c, insulin and glucose are a combination of measures from cord blood, fasting, non-fasting bloods, and NMR. Interleukin-6 (IL-6) was measured by ELISA (R&D systems, Abingdon, UK) and available at nine years, and C-reactive protein (CRP) was measured by automated particle-enhanced immunoturbidimetric assay (Roche UK, Welwyn Garden City, UK). CRP levels were measured at nine, 15 and 18 years of age. More details of measurement assays and protocols are provided in Supplementary material.

#### Physical Activity

All children who attended the clinic at ages 11, 13, and 15 years were asked to wear an accelerometer for 7 days. More details of the physical activity measures are provided in Supplementary material.

##### Genetic data

DNA samples from 9,912 ALSPAC children were genotyped on the Illumina HumanHap550-quad SNP array genotyping platform. After quality control assessment methods, imputation, and restriction to 1 child per family, genetic data was available for 7,977 individuals. More details are in Supplementary material.

## STATISTICAL ANALYSIS

### Generating polygenic risk scores

PRS were computed according to the method described by the International Schizophrenia Consortium [17], based on the summary statistics from the genome-wide association study (GWAS) of Alzheimer’s disease by the IGAP consortium [8]. Details about IGAP are in the Supplementary material. PRS were created for the significance thresholds: 5×10^−8^ (genome-wide significance threshold), 5×10^−2^, and 5×10^−1^, including a different number of SNPs (Supplementary table 1). A list of the 17 SNPs used to generate the PRS at the genome-wide significant threshold is provided in Supplementary table 2. This threshold was chosen for the main analysis since it includes the most strongly associated SNPs. However, less stringent p-value thresholds can explain more of the Alzheimer’s disease variance. Results for the other thresholds are in Supplementary material. As *ApoE i*s involved in lipid homeostasis by mediating lipid transport [18], the main analysis used scores excluding *ApoE.*

### Modelling trajectories of cardiometabolic risk factors

Children with at least one measure were included in the multilevel models to minimise selection bias, under a missing at random assumption [19, 20]. All trajectories were estimated using linear spline multilevel models (2 levels: measurement occasion and individual). The optimal linear spline model for each cardiometabolic risk factor was based on previous work [20-23]. All trajectories were modelled in MLwiN version 3.01 [24], called from STATA version 14 [25] using the runmlwin command [26]. All trajectories were centred on the lowest age (in whole years) at the first available measure and mean PRS. Variables with a skewed distribution (triglycerides, fat mass, CRP) were (natural) log transformed. Fat and lean mass were adjusted for height with the use of a time- and sex-varying height covariate derived in previous literature [27], which was centred on the mean of this derived covariate. The non-linearity of height trajectories was modelled using fractional polynomials to find the best fitting function of age. Further details of trajectories are in Supplementary material. We incorporated a fixed effect for the PRS in each trajectory along with an interaction term for the PRS and the linear splines or fractional polynomials. The coefficients represent the mean difference in the risk factor at baseline per one standard deviation (SD) higher PRS, and the mean change in the risk factor per year, or per year within each linear spline period, per one SD higher PRS. Results including the logged risk factors are interpreted as mean percentage change rather than mean difference. The mean trajectory was allowed to differ between males and females by including an interaction between gender and age. Results were interpreted according to the American Statistical Association guidance [28, 29].

#### Modelling risk factors with sparse measures

Due to the sparse measures for birthweight, insulin, IL-6, and MVPA, linear regression models were used to estimate the association between the PRS and these measures. Measures from the clinic with the highest number of participants were used for the analysis, where applicable. As insulin, IL-6 and MVPA had a skewed distribution, they were transformed using the natural logarithm. Models were adjusted for age, sex and the first three ancestry-informative principal components.

##### Sensitivity analyses

We examined the associations between the PRS including only the two SNPs tagging the *ApoE* region (rs7412 and rs429358) and the PRS excluding *ApoE* and cardiometabolic risk factors at different p-value thresholds. To examine if our results were changed by the inclusion of non-fasted bloods in some of the clinics, we repeated analyses excluding all participants who reported eating four hours prior to the 15 and 18-year clinics. To test the generalisability of our results, we investigated the characteristics of individuals in the analysis of insulin (model with the fewest individuals and repeated measures) to those excluded from this analysis (individuals with no genetic or cardiometabolic data).

## RESULTS

### Descriptive statistics

The number of children used in the models ranged from 699 to 6,953 individuals. Descriptive and model fit statistics are in Supplementary tables S3-4. The household social class of children in the analysis of glucose (model with the smallest sample size) was higher compared to children excluded due to missing data (19.6% professional vs. 12.3% professional, p<0.0001). Furthermore, the prevalence of maternal smoking during pregnancy was lower in the included sample compared to the excluded sample (17.5% vs 28.2%, p<0.0001) (Supplementary table S5).

### Main analysis

Figures 1-3 show the mean trajectories of anthropometry, blood pressure, and blood-based biomarkers measures, respectively, alongside the association between the PRS and each of these cardiometabolic risk factor trajectories. At the genome-wide significant p-value threshold, there was weak evidence of association of a PRS with birthweight (Supplementary table S6.1) and height trajectories (Figure 1A). At age nine years, there was weak evidence that a PRS was associated with a higher height-adjusted fat mass (β: 0.59%; 95% confidence interval [CI]: −0.92, 2.11, Supplementary table S6.3) and height-adjusted lean mass (β: 0.04 kg; 95% CI: −0.03, 0.11, Supplementary table S6.3). In line with these results, there was little evidence of an association between the PRS, SBP, glucose, triglycerides, non-HDL, HDL-c and CRP (Figure 3). There was stronger evidence of an association with higher DBP (β: 0.19 mmHg; 95% CI: 0.02, 0.37) at age 7 years and lower DBP (β: −0.05 mmHg; 95% CI: −0.09, - 0.003) from ages 7 to 12 years (Table S6.4). Cross-sectional models showed weak evidence of association with higher insulin, IL-6, and MVPA levels (Supplementary table S6.8-6.9). The direction of effect between time periods in multilevel models was consistent only for CRP (Supplementary table S6.7) and MVPA (Supplementary table S6.9). For all risk factors except for DBP, the 95% CIs spanned the null.

**Figure 1.**
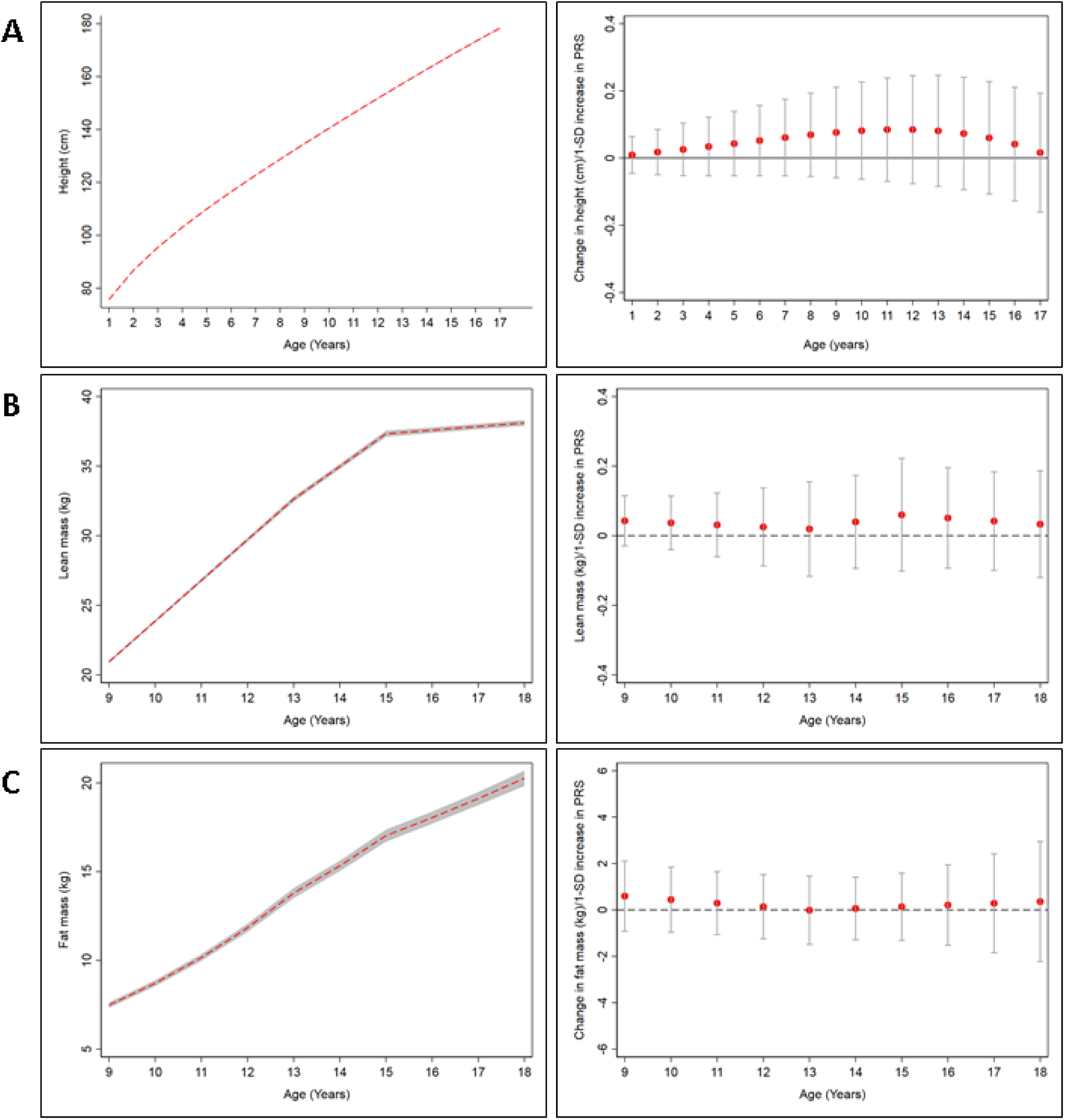
Mean trajectories of anthropometric measures and mean change in anthropometric measures per 1-SD increase in PRS from ages 9-18 years in ALSPAC. (A)height (B) lean mass, and (C)fat mass. The shaded areas in the left graphs and the bars in the right graphs represent 95% confidence intervals, respectively.

**Figure 2.**
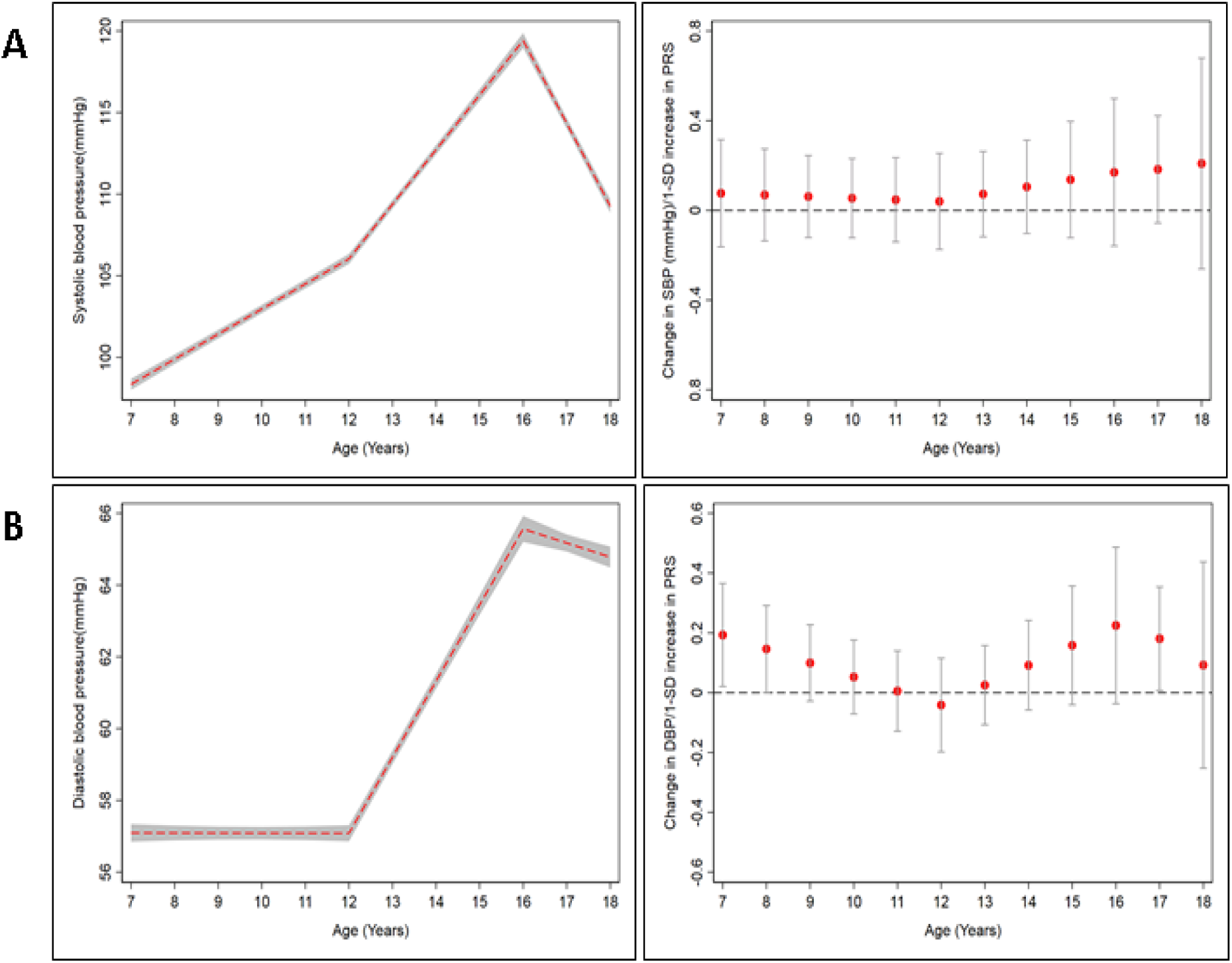
Mean trajectories of blood pressure measures and mean change in blood pressure measures per 1-SD increase in PRS from ages 7-18 years in ALSPAC. (A) systolic blood pressure, (B) diastolic blood pressure. The shaded areas in the left graphs and the bars in the right graphs represent 95% confidence intervals, respectively.

**Figure 3.**
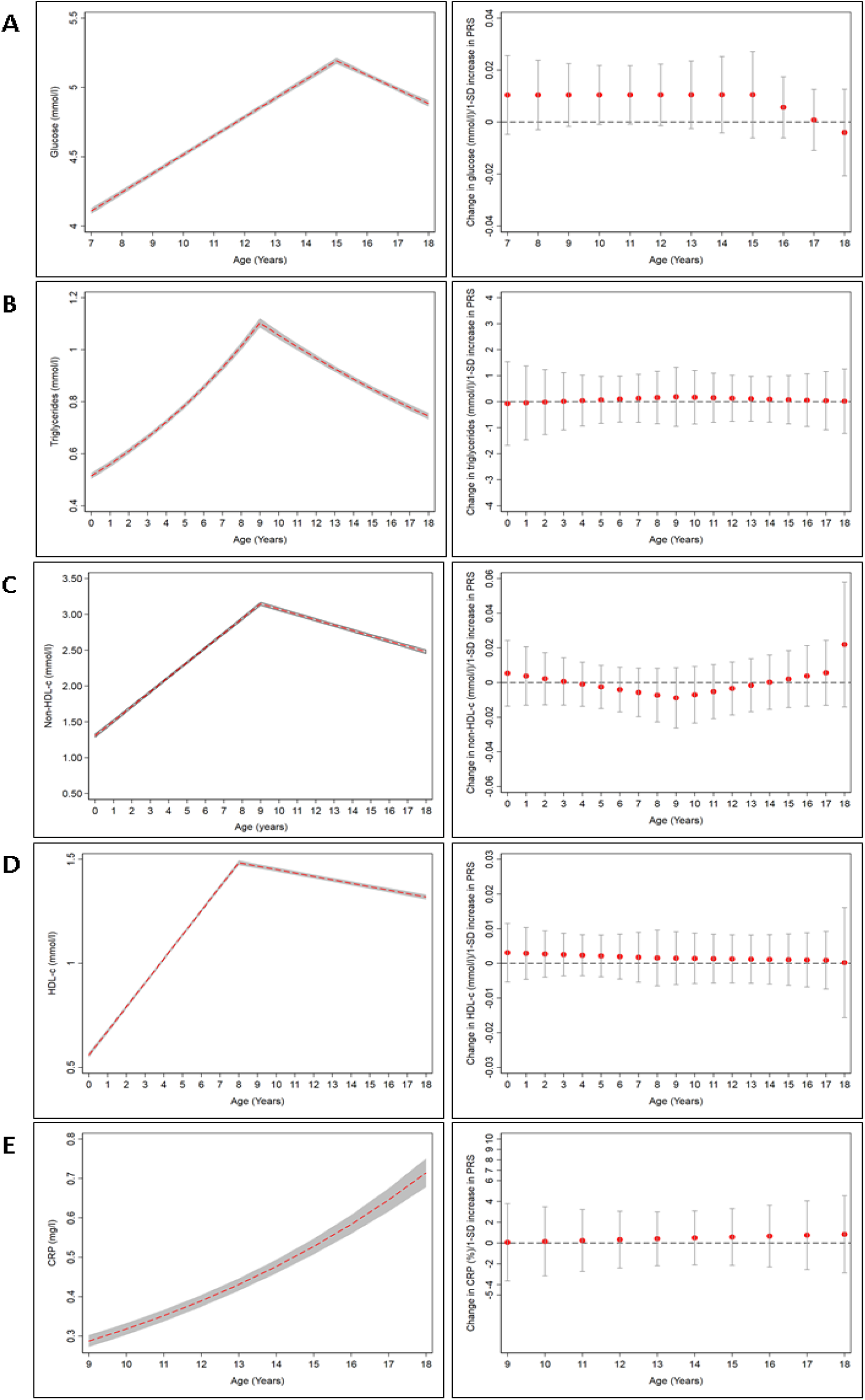
Mean trajectories of blood-based biomarkers and mean change per 1-SD increase in PRS. (A) glucose from ages 7-18 years, (B) triglycerides, (C) non-HDL-c, and (D) HDL-c from ages 0-18 years, (E) CRP from ages 9-18 years. The shaded areas in the left graphs and the bars in the right graphs represent 95% confidence intervals, respectively.

### Sensitivity analyses

Sensitivity analyses comparing the results for the PRS at different p-value thresholds used PRS excluding *ApoE*. Further analyses comparing the results with the PRS including/excluding *ApoE* were performed using the genome-wide significant threshold. The effect size for the β-coefficients was larger for risk factors such as height, lean mass, triglycerides, CRP at the more liberal PRS thresholds. A higher PRS was associated with shorter stature at the more liberal thresholds at ages 4-10 years (β: - 0.15 cm; 95% CI: −0.20, −0.01 at age 5 years and p≤5×10^−1^, Supplementary table S8.2). A higher PRS was associated with lower lean mass at ages 9 and 15-18 years (β: −0.06kg/yr; 95% CI: −0.11, −0.02 at age 15 years and p≤5×10^−1^, Supplementary table S8.3) The direction of effect differed between the genome-wide significant threshold and the more liberal thresholds for some risk factors (Supplementary tables S7-10). PRS which incorporated the *ApoE* region were associated with cardiometabolic risk factors such as HDL-c, non-HDL-c, DBP, and CRP as expected by the role of *ApoE* in lipid homeostasis. There was strong evidence of association between a PRS including the *ApoE* region and higher non-HDL-c levels at birth (β: 0.03; 95% CI: 0.01, 0.04, Supplementary table S9.6) and from ages 9-12 years (β: −0.003; 95% CI: −0.01, −0.0005). There was also strong evidence that a PRS with *ApoE* was associated with lower HDL-c levels at birth (β: −0.01 mmol/l; 95% CI: - 0.02, −0.005, Supplementary table S9.6) and from birth to age 7 years (β: −0.002 mmol/l yr; 95% CI: - 0.003, −0.0004), and higher HDL-c levels from ages 7-18 years (β: 0.001 mmol/l yr; 95% CI: 0.0002, 0.002). For a 1 SD increase in the PRS for, CRP levels were 8.60% (95% CI; −11.86, −5.21, Supplementary table S9.7) lower at age 9 years. To compare fasting/nonfasting biomarkers, we used the PRS excluding *ApoE* at the genome-wide significant threshold. The associations observed between non-fasted and fasted bloods were similar for all examined blood-based biomarkers (Supplementary tables S10).

## DISCUSSSION

This is the first study examining whether a PRS for Alzheimer’s disease was associated with trajectories of cardiometabolic risk factors through childhood and adolescence. We examined these associations firstly to test whether associations may emerge early in the life course, and secondly to minimise selection bias due to attrition from cardiometabolic-associated morbidity and mortality. We did not find any consistent evidence to suggest that the PRS for Alzheimer’s disease was associated with cardiometabolic risk factors in childhood or adolescence. The predominantly null results at the genome-wide significant p-value threshold may indicate that either the PRS for Alzheimer’s disease does not operate through cardiometabolic pathways or that it operates partly through cardiometabolic pathways, but the associations emerge much later in life.

In our sensitivity analyses, we observed associations between the more liberal PRS and some of the cardiometabolic risk factors (such as height, lean mass, triglycerides, insulin, and CRP). For example, we found evidence of association with shorter stature from ages 4-10 years, lower lean mass at ages 9 years and from ages 15-18 years (at p-value threshold p≤5×10^−1^). As these PRS include many genetic variants which are weakly associated with Alzheimer’s disease and the associations observed are dissimilar in the direction of effect for all time-points, there is a chance that the associations may be attributed to horizontal pleiotropy where the PRS is associated with a range of independent phenotypes. Furthermore, the effects of the PRS at each p-value threshold cannot be disentangled since PRS at higher p-value thresholds include all SNPs from PRS at lower p-value thresholds. We also found evidence of an association between the PRS for Alzheimer’s disease (including the *ApoE* region) and HDL, non-HDL, CRP, and triglyceride levels, but the direction of effect was not consistent across all time-points.

### Comparison to other studies

Longitudinal cohort studies with prospective phenotypic data through early-life into adulthood and clinical diagnosis of Alzheimer’s disease are scarce. Studies researching developmental factors such as height are the only source of data on early life exposures. It has been hypothesised that suboptimal foetal development, indexed by small birth weight/size may result in permanent changes in the structure, metabolism of the organs through various biological mechanisms [30]. Contrary, positive associations have been found between several cardiometabolic risk factors in midlife and Alzheimer’s disease [1], with higher DBP and SBP increasing the risk of Alzheimer’s disease [5], independent of the *ApoE* genotype. A positive association has also been found for BMI trajectories in midlife and Alzheimer’s disease [31].

A study of PRS for Alzheimer’s disease in adults with Alzheimer’s disease and/or a family history of Alzheimer’s disease identified an inverse association between a PRS, hypertension and stroke, but no associations for type 2 diabetes and heart disease [32]. Studies in children have only focused on *ApoE*, where there is an established linear relationship between *ApoE* and LDL in children and adults [33, 34]. Although the *ApoE* region is the largest genetic risk factor for Alzheimer’s disease, it is also strongly independently associated with cardiometabolic risk factors (primarily lipids), making it difficult to exclude pleiotropy as an explanation for any observed findings (i.e. *ApoE* may be associated with cardiometabolic risk independently of Alzheimer’s disease. Contrary to a recent GWAS [35], we did not find an association between *ApoE4* and physical activity. In agreement with our results, a study found a negative genetic correlation between height and dementia, suggesting our findings for height at liberal thresholds may be due to shared genetic pathways [36].

### Strengths and limitations

We performed our analysis in children and adolescents to examine whether genetic risk for Alzheimer’s disease operates through cardiometabolic pathways in early in the life course and to minimise selection bias due to attrition from cardiometabolic-associated morbidity and mortality. Additionally, the use of genetic scores makes confounding by social and lifestyle characteristics highly unlikely. Our study benefits from a well-characterised cohort with repeated measures of a range of cardiometabolic risk factors over a course of 18 years. The use of multilevel models enabled us to account for the clustering of repeated measurements within individuals and the correlation between repeated measurements over time.

However, as genetic scores explain a small proportion of variance in the outcome, it is highly likely we may have been underpowered to detect small polygenic effects on the examined cardiometabolic risk factors. For this reason, we did not correct our results for multiple testing, as our findings were largely null. Further limitations include the use of non-fasting and fasting bloods for some of the examined risk factors and the availability of measures from birth for only four of the risk factors. However, the estimates for fasted and non-fasted bloods were similar in our sensitivity analyses. Furthermore, PRS were derived from a GWAS of European individuals and our study was also conducted in individuals of European ancestry; hence, these results may not be generalizable to other populations.

## Conclusion

We found little evidence to suggest that the combined genetic effects conferring an increased risk for Alzheimer’s disease were associated with cardiometabolic risk factors, beyond the ApoE effect. As this is the first study to examine these associations through childhood and adolescence, these findings should be replicated in other large birth cohorts to examine whether the genetic risk for Alzheimer’s disease can be captured in early childhood, or whether it becomes phenotypically manifest in adulthood.

## Acknowledgements

We are extremely grateful to all the families who took part in this study, the midwives for their help in recruiting them, and the whole ALSPAC team, which includes interviewers, computer and laboratory technicians, clerical workers, research scientists, volunteers, managers, receptionists and nurses.

We also thank the International Genomics of Alzheimer’s Project (IGAP) for providing summary results data for these analyses. The investigators within IGAP contributed to the design and implementation of IGAP and/or provided data but did not participate in analysis or writing of this report. IGAP was made possible by the generous participation of the control subjects, the patients, and their families. The i–Select chips was funded by the French National Foundation on Alzheimer’s disease and related disorders. EADI was supported by the LABEX (laboratory of excellence program investment for the future) DISTALZ grant, Inserm, Institut Pasteur de Lille, Université de Lille 2 and the Lille University Hospital. GERAD was supported by the Medical Research Council (Grant n° 503480), Alzheimer’s Research UK (Grant n° 503176), the Wellcome Trust (Grant n° 082604/2/07/Z) and German Federal Ministry of Education and Research (BMBF): Competence Network Dementia (CND) grant n° 01GI0102, 01GI0711, 01GI0420. CHARGE was partly supported by the NIH/NIA grant R01 AG033193 and the NIA AG081220 and AGES contract N01–AG–12100, the NHLBI grant R01 HL105756, the Icelandic Heart Association, and the Erasmus Medical Center and Erasmus University. ADGC was supported by the NIH/NIA grants: U01 AG032984, U24 AG021886, U01 AG016976, and the Alzheimer’s Association grant ADGC–10–196728.

