## Supplementary Material for "Polygenic risk score for Alzheimer’s disease and trajectories of cardiometabolic risk factors in children"

**Methods**

**Measures**

#### **Genetic data**

A total of 9,912 ALSPAC children were genotyped on the Illumina HumanHap550-quad SNP genotyping platform. The resulting raw genome-wide data were subjected to standard quality control methods. Individuals were excluded on the basis of gender mismatches; minimal or excessive heterozygosity; disproportionate levels of individual missingness (>3%) and insufficient sample replication (IBD < 0.8). Population stratification was assessed by multidimensional scaling analysis and compared with Hapmap II (release 22) European descent (CEU), Han Chinese, Japanese and Yoruba reference populations; all individuals with non-European ancestry were removed. SNPs with a minor allele frequency of <1%, a call rate of < 95% or evidence for violations of Hardy-Weinberg equilibrium (P < 5x10^-7^) were removed. Cryptic relatedness was measured as proportion of identity by descent (IBD > 0.1). Related subjects that passed all other quality control thresholds were retained during subsequent phasing and imputation. Following quality control procedures, imputation, exclusion of individuals with withdrawn consent and restricting to 1 child per family, genetic data was available for 7,977 children.

**Discovery sample**

The dataset reported by the International Genomics of Alzheimer's Project (IGAP) consortium [2], consisting of 17,008 Alzheimer's disease cases and 37,154 controls was used as the discovery sample. IGAP used genotyped and imputed data on 7,055,881 single nucleotide polymorphisms (SNPs) to meta-analyse four previously-published GWAS datasets consisting of EADI (European Alzheimer's Disease Initiative), ADGC (Alzheimer Disease Genetics Consortium), CHARGE (Cohorts for Heart and Aging Research in Genomic Epidemiology), and GERAD (Genetic and Environmental Risk in AD). Complete details of each study, as well as the samples and methodologies are reported elsewhere [2–4]. Each dataset was imputed with either Impute 2 [5] or MACH software [6], utilising the 1000 genomes data as a reference panel.

**Polygenic risk score (PRS)**

SNPs were removed from the analysis if there was an allelic mismatch between samples (the alleles reported by the IGAP study did not match the alleles present in the ALSPAC sample). Correlated SNPs were removed by a pruning procedure in PLINK genetic analysis tool [7] (--clump command using an r^2^ parameter of 0.25 and a physical distance threshold for clumping SNPs of 500 kB). A PRS was calculated for each participant with genetic data using PLINK (version 1.9). Each score was calculated from the effect size (logarithm (log)odds))-weighted sum of associated alleles within each participant). SNPs at the CLU (rs9331896) and the HLA region (rs9271192) were not present in ALSPAC and were omitted from the PRS. The PRS was standardised by subtracting the mean and dividing by the standard deviation.

**Phenotypic measures**

*Description of measurement sources*

*Details on measurement of systolic (SBP) and diastolic blood pressure (DBP)*

A Dinamap 9301 Vital Signs Monitor (Morton Medical, London) was used at the seven, nine, 9, and 11-year clinics, 15- and 18-year clinics; an Omron MI-5 was used at the 10-year clinic; a Dinamap 8100 Vital Signs Monitor (Morton Medical) was used at the 13-year clinic.

*Details on measurement of blood-based biomarkers*

Plasma lipid assays (total cholesterol, triglycerides, and high-density lipoprotein cholesterol (HDL-c) were conducted by modification of the standard Lipid Research Clinics Protocol using enzymatic reagents for lipid determination. Samples were collected after an overnight fast and were analysed by the hexokinase method. High-sensitivity interleukin-6 (IL-6) was measured by enzyme-linked immunosorbent assay (R&D systems, Abingdon, UK) and CRP was measured by automated particle-enhanced immunoturbidimetric assay (Roche UK). Insulin was measured by an ELISA (Mercodia, Uppsala, Sweden), which does not cross-react with proinsulin. All assay coefficients of variation were <5%. Non-fasting blood samples were obtained using standard procedures with samples immediately spun and frozen at -80ºC. The measurements were assayed after a median of 7.5 years in storage with no prior freeze–thaw cycles during this period.

*Measurement of biomarkers using* nuclear magnetic resonance *(NMR) spectroscopy*

A comprehensive profiling of offspring circulating lipids, lipoproteins, and metabolites was performed by a high-throughput NMR metabolomics platform, providing a snapshot of offspring serum metabolome at follow-up [8,9]. At age seven, this was performed on fasted blood samples and glucose is included in our analyses. At ages 15 and 18 years, non-fasted bloods were used.

*Measurement of MVPA using actigraphy*

All children who attended the clinic at ages 11, 13, and 15 years were asked to wear an Actigraph AM7164 2.2 accelerometer (Actigraph LLC, Fort Walton Beach, FL, USA) around their waist, at the right hip for 7 days. Average minutes of moderate to vigorous physical activity (MVPA) per valid day was defined as a value of greater than 3600 counts per minute, based on a previous calibration study [1]. Data were considered valid only if children had worn the accelerometer for at least 10 hours a day for at least 3 days. Ten or more minutes of successive zeros were regarded as non-wear time and were deleted from each file.

**Details of model selection**

Details of model selection have been published elsewhere [12–16]. A brief description is provided below.

**Height** was modelled by including non-linear age functions in a multilevel model. Models incorporated powers of age (0,0.5,2,3, where a power of zero is the log function), and each combination of pairs of these powers.

**Height-adjusted fat mass and height-adjusted lean mass** were measured on five occasions between 9 and 18 years. Knots were placed at 13, 15 and 18 resulting in three periods of change; from 9-13, 13-15, 15-18. Both models were adjusted for a time- and sex-varying height covariate which was incorporated as a fixed effect. The variance of measurement occasion-level residuals (the differences between observed and predicted measurements) was allowed to differ with age. The models assumed the form of: height-adjusted fat mass_ij_/height-adjusted lean mass_ij_= β_0_ +u_0j_ +(β_1_+ u_1j_)s_ij1_ +(β_2_+ u_2j_ )s_ij2_ + (β_3_ +u_3j_) _Sij3_+β_4_(PRS)+β_5_(PRS)s_ij1_+β_6_(PRS)s_ij2_+β_7_ (PRS)s_ij3_+β_8_(sex)+β_9_(sex)s_ij1_+β_10_(sex)s_ij2_+β_11_(sex) s_ij3_+β_12_(PC1)+β_13_(PC2)+β_14_(PC3)+β_15_(age and sex-adjusted height covariate)_ij_ + e_ij_(age_binary_ij_) where for individual j at measurement occasion i; β_0_ is the fixed effect coefficient for the average intercept, β_1_-β_3_ represent fixed effect coefficients for the average linear slopes of each linear spline, β_4_ represents the fixed effect coefficient for the association between the PRS for Alzheimer’s disease and the average intercept, β_5_-β_7_ represent the fixed effect coefficients for the association of a 1 SD increase in the PRS with change per year in the outcome in each spline period, β_8_ represents the fixed coefficient for the association of sex with the average intercept and β_9_-β_11_ are the fixed coefficients for the association of sex with the average slope of each linear spline, β_12_-β_14_ represent the fixed coefficients for the principal components, β_15_ represents the fixed effect coefficient for the average difference in measurements between individuals of different heights, S, u_0_-u_3_ are individual-specific random effects for the intercept and slopes respectively, and e_ij_ represents the occasion-specific residuals or measurement error which was allowed to change with age>13 and age<14.

**SBP** and **DBP** were measured at seven time-points from 7 to 18 years. The knots for both models were placed at 12, 16, and 18 resulting in three periods of change; from seven to 12, 12-16 and 16-18. Both models included a fixed effect to control for the use of a different machine (Omron MI-5 machine) for measuring blood pressure in 10-year clinic and a binary time indicator as a level one random effect of age less than or greater than 10 years to account for varying measurement error with age. The models took the form of: SBP_ij_ = β_0_ + u_0j_ + (β_1_+ u_1j_)s_ij1_ + (β_2_+ u_2j_ )s_ij2_ + (β_3_ + u_3j_) _Sij3_+ β_4_(PRS) +β_5_(PRS)s_ij1_+β_6_ (PRS)s_ij2_+ β_7_(PRS)s_ij3_+ β_8_(sex)+ β_9_(sex) s_ij1_+β_10_(sex) s_ij2_+ β_11_(sex) s_ij3_+ + β_12_(PC1)+ β_13_(PC2)+ β_14_(PC3)+ β_15_(machine)+e_ij_(age_binary_ij_)**,** where for individual j at measurement occasion i; β_0_ is the fixed effect coefficient for the average intercept, β_1_-β_3_ represent fixed effect coefficients for the average linear slopes of each linear spline, β_4_ represents the fixed effect coefficient for the association between the PRS and the average intercept, β_5_-β_7_ represent the fixed effect coefficients for the association of a 1 SD increase in the PRS with change per year in the outcome in each spline period , β_8_ represents the fixed coefficient for the association of sex with the average intercept and β_9-_β_11_ are the fixed coefficients for the association of sex with average slope of each linear spline, β_12_-β_14_ represent the fixed coefficients for the principal components,β_13_ represents the fixed effect for the average difference in measurements between the machine used at the 10-year clinic compared to the machine used at other clinics, u_0_-u_3_ indicate person-specific random effects for the intercept and slopes respectively, and e_ij_ is the occasion-specific residuals or measurement error which was allowed to vary with age.

**Triglycerides** and **HDL-c** were measured five times from birth to 18 years. Non-HDL-c was obtained by deducting HDL-c from total cholesterol. Knots for triglycerides and non-HDL were set at 9 and 18 years resulting in two periods of change; from birth to 9 years and 9 to 18 years. Knots for HDL were placed at 8 and 18 years resulting in two periods of change; from birth to 8 years and 8 to 18 years. The models for triglycerides/HDL-c/non-HDL-c took the form of: triglycerides_ij_/HDL-c_ij_/non-HDL-c_ij_ = β_0_ + u_0j_ + (β_1_+ u_1j_)s_ij1_ + (β_2_+ u_2j_ )s_ij2_ + β_3_(PRS) +β_4_ (PRS)s_ij1_+ β_5_ (PRS)s_ij2_+ β_6_(sex)+ β_7_(sex) s_ij1_+β_8_(sex) s_ij2_+ β_9_(PC1)+ β_10_(PC2)+ β_11_(PC3) + e_ij_ where for person j at measurement occasion i; β_0_ is the fixed effect coefficient for the average intercept, β_1_-β_2_ represent fixed effect coefficients for the average linear slopes of each linear spline, β_3_ represents the fixed effect coefficient for the association between the PRS and the average intercept, β_4_-β_5_ represent the fixed effect coefficients for the association of a 1 SD increase in the PRS with change per year in the outcome in each spline period, β_6_ represents the fixed coefficient for the association of sex with the average intercept and β_7-_β_8_ are the fixed coefficients for the association of sex with average slope of each linear spline, β_9_-β_11_ represent the fixed coefficients for the principal components,u_0_-u_3_ indicate person-specific random effects for the intercept and slopes respectively, and e_ij_ represents the occasion-specific residuals or measurement error.

**Glucose** was measured on four occasions (7, 9, 15, and 18). Knots were placed at 15 and at 18 resulting in two periods of change; from seven to 15 and 15-18. Due to sparse number of available measurements of glucose, we modelled the person-specific random effects as a single linear slope instead of a function of the splines as has been done in other linear models. This allowed for individual-specific variation from the mean trajectory but under the assumption that individual-specific variation was constant over time. The model took the form of: glucose_ij_ = β_0_ + u_0j_ + (β_1_)s_ij1_ + (β_2_)s_ij2_ + β_3_(PRS)+β_4_(PRS)s_ij1_+ β_5_(PRS)s_ij2_ + β_6_(sex)+ β_7_(sex)s_ij1_+β_8_(sex)s_ij2_+β_9_(PC1)+ β_10_(PC2)+ β_11_(PC3) + u_1j_*age + e_ij_ where for person j at measurement occasion i; β_0_ is the fixed effect coefficient for the average intercept, β_1_-β_2_ represent fixed effect coefficients for the average linear slopes of each linear spline, β_3_ represents the fixed effect coefficient for the association between the PRS and the average intercept, β_4_-β_5_ represent the fixed effect coefficients for the association of a 1 SD increase in the PRS with change per year in the outcome in each spline period , β_6_ represents the fixed coefficients for the association of sex with the average intercept and β_7-_β_8_ are the fixed coefficients for the association of sex with average slope of each linear spline, β_9_-β_11_ represent the fixed coefficients for the principal components, u_0j_ to u_1j_ indicate person-specific random effects for the intercept and slope respectively, and e_ij_ represents the occasion-specific residuals or measurement error.

**Details of adjustment for height in models of height-adjusted lean mass and height-adjusted fat mass**

Analyses of adiposity usually adjust for height squared to eliminate the effect of height, allowing for the residual to represent adiposity which is then assumed to be independent of height, since taller height is associated with higher weight or overall mass. As height varies substantially across childhood and adolescence, suitable age and sex-specific adjustment for height is needed to accurately measure sex differences in height-adjusted lean mass or height-adjusted fat mass over time which are independent of height. In an analysis in the ALSPAC cohort, with several repeated measures of BMI from birth to 16 years, powers of height adjustment differed over time. Around puberty, sex differences in this were also observed where BMI was adjusted for height to the power of 2.5 in males compared to height to the power of 1.8 in females suggesting that many existing studies of anthropometry in youth could potentially overestimate or underestimate BMI among the sexes during different age periods due to their differential heights [10]. We used appropriate powers of height adjustment which were age and sex-specific for inclusion in multilevel models of height-adjusted lean mass and fat mass which were derived and reported elsewhere [11,12].

**Variables used to compare the socioeconomic and health traits of mothers of children in the inclusion sample compared to the exclusion sample for the insulin model**

We investigated the characteristics associated with being excluded from our analyses due to missing data on the risk factor of interest or genetic data. We compared the socio-demographic characteristics at birth of mothers and partners of children included in the analysis of insulin compared to those excluded from the analysis. Insulin was chosen as it had the fewest repeated measures over time. All characteristics were measured during pregnancy or at birth through questionnaires or from routine health records. Marital status was acquired from antenatal questionnaires and classified as never married, widowed, divorced, separated, first marriage, marriage 2 or 3. Household social class was measured as the highest of the mother’s or her partner’s occupational social class using data on job title and occupation collected about the mother and her partner from the mother’s questionnaire at 32 weeks of gestation. Social class was derived using the standard occupational classification (SOC) codes developed by the United Kingdom Office of Population Census and Surveys and classified as I professional, II managerial and technical, IIINM non-manual, IIIM manual, and IV&V part skilled occupations and unskilled occupations. A questionnaire at 32 weeks of gestation, mothers were asked to report their educational attainment, which was categorized as below O-Level (Ordinary Level; exams taken in different subjects usually at age 15-16 at the completion of legally required school attendance, equivalent to today’s UK General Certificate of Secondary Education), O-Level only, A-Level (Advanced-Level; exams taken in different subjects usually at age 18), or university degree or above. For a questionnaire at 32 weeks of gestation, partners were asked to report their educational attainment, which was categorized as below O-Level, O-Level only, A-Level, or university degree or above. Smoking in the first trimester of pregnancy was self-reported by mothers at 18 weeks of gestation; responses to smoking any tobacco (cigarettes, cigars, pipes, or other) were categorised accordingly: no smoking, <10 per day, 10-19 per day or greater than 19 per day. Clinical records were used to estimate gestational age at birth.

| **Table S1** Number of SNPs included in PRS at different p-value thresholds including/excluding the ApoE region | | |
| --- | --- | --- |
| **P-value threshold** | **ApoE** | **No ApoE** |
| 5x10^-8^ | 19 | 17 |
| 5x10^-2^ | 44,893 | 45,040 |
| 5x10^-1^ | 240,803 | 240,561 |
| ApoE; apolipoprotein-E; PRS, polygenic risk scores; SNPs, single nucleotide polymorphisms | | |

GWAS, genome-wide association study; SNPs, single nucleotide polymorphisms.

^a^SNPs included in the genetic risk score (GRS). Additional information provided: chromosomal and base pair position, minor allele (A1) and major allele (A2), odds ratio (OR).

| **Table S2** Genome-wide significant SNPs included in the genetic risk score based on IGAP GWAS Stage 1[19] | | | | | | |
| --- | --- | --- | --- | --- | --- | --- |
| SNP^a^ | Chromosome | Position | Nearest gene | A1 | A2 | OR |
| rs6656401 | 1 | 207,692,049 | CR1 | A | G | 1.17 |
| rs6733839 | 2 | 127,892,810 | BIN1 | T | C | 1.21 |
| rs35349669 | 2 | 234,068,476 | INPP5D | T | C | 1.07 |
| rs190982 | 5 | 88,223,420 | MEF2C | G | A | 0.92 |
| rs10948363 | 6 | 47,487,762 | CD2AP | G | A | 1.10 |
| rs2718058 | 7 | 37,841,534 | NME8 | G | A | 0.93 |
| rs1476679 | 7 | 100,004,446 | ZCWPW1 | C | T | 0.92 |
| rs11771145 | 7 | 143,110,762 | EPHA1 | A | G | 0.90 |
| rs28834970 | 8 | 27,195,121 | PTK2B | C | T | 1.10 |
| rs10838725 | 11 | 47,557,871 | CELF1 | C | T | 1.08 |
| rs983392 | 11 | 59,923,508 | MS4A6A | G | A | 0.90 |
| rs10792832 | 11 | 85,867,875 | PICALM | A | G | 0.88 |
| rs11218343 | 11 | 121,435,587 | SORL1 | C | T | 0.76 |
| rs17125944 | 14 | 53,400,629 | FERMT2 | C | T | 1.13 |
| rs10498633 | 14 | 92,926,952 | SLC24A4 | T | G | 0.90 |
| rs4147929 | 19 | 1,063,443 | ABCA7 | A | G | 1.14 |
| rs7274581 | 20 | 55,018,260 | CASS4 | C | T | 0.87 |

Table S3.1 Descriptive statistics of height and PRS by inclusion and exclusion criteria

|  | INCLUDED SAMPLE | | EXCLUDED SAMPLE | |
| --- | --- | --- | --- | --- |
| Outcome | **Participants (N)** | **Mean (cm) (SD)** | **Participants (N)** | **Mean (cm) (SD)** |
| Height |  |  |  |  |
| 1 years | 5,379 | 81.19 (4.56) | 9,526 | 81.24 (4.52) |
| 5 years | 3,565 | 114.28 (5.99) | 5,204 | 114.16 (6.11) |
| 10 years | 5,375 | 143.14 (6.59) | 7,449 | 143.03 (6.63) |
| 18 years | 2,823 | 171.47 (9.31) | 10,436 | 171.11 (9.32) |
| PRS | 7,844 | -0.0002 (1.00) | 133 | 0.02 (1.00) |

cm, centimetres; PRS; polygenic risk score; SD, standard deviation.

Table S3.2 Descriptive statistics of height-adjusted fat, lean mass and PRS by inclusion and exclusion criteria

|  | INCLUDED SAMPLE | | EXCLUDED SAMPLE | |
| --- | --- | --- | --- | --- |
| Outcome | **Participants (N)** | **Mean (kg) (SD)** | **Participants (N)** | **Mean (kg) (SD)** |
| Height-adjusted fat mass |  |  |  |  |
| 9 years | 5,300 | 8.45 (4.91) | 1,964 | 8.64 (5.21) |
| 9-13 years | 5,107 | 11.57 (6.51) | 1,935 | 11.96 (7.08) |
| 13-15 years | 4,491 | 13.69 (7.83) | 1,567 | 13.70 (8.11) |
| 15-18 years | 4,458 | 16.55 (9.54) | 1,968 | 16.90 (9.97) |
| PRS | 6,181 | 0.002 (1.00) | 1,678 | -0.01 (1.00) |
| Height-adjusted lean mass |  | | | |
| 9 years | 5,309 | 24.58 (3.17) | 1,974 | 24.54 (3.35) |
| 9-13 years | 5,115 | 29.75 (4.31) | 1,852 | 29.75 (4.54) |
| 13-15 years | 4,500 | 38.04 (6.42) | 1,512 | 37.73 (6.47) |
| >15 years | 4,468 | 44.39 (9.27) | 1,739 | 44.03 (9.33) |
| PRS | 6,188 | 0.002 (1.00) | 1,671 | -0.005 (1.00) |

kg, kilogram; PRS; polygenic risk score; SD, standard deviation.

**Table S3.3** Descriptive statistics for SBP and DBP, PRS by inclusion and exclusion criteria

|  | INCLUDED SAMPLE | | EXCLUDED SAMPLE | |
| --- | --- | --- | --- | --- |
| Outcome | **Participants (N)** | **Mean (mmHg) (SD)** | **Participants (N)** | **Mean**  **(mmHg) (SD)** |
| SBP |  |  |  |  |
| 7 years | 5,745 | 98.84 (9.12) | 2,330 | 99.25 (9.29) |
| 7-12 years | 6,007 | 103.93 (9.33) | 2,901 | 103.00 (9.74) |
| 12-16 years | 5,421 | 112.76 (11.90) | 2,044 | 112.61 (11.98) |
| 16-18 years | 3,505 | 115.08 (10.20) | 1,398 | 115.21 (10.48) |
| PRS | 6,733 | -0.003 (1.00) | 1,126 | 0.02 (1.00) |
| DBP |  |  |  |  |
| 7 years | 5,743 | 56.30 (6.59) | 2,331 | 56.93 (6.74) |
| 7-12 years | 6,008 | 58.61 (7.01) | 2,799 | 59.19 (7.19) |
| 12-16 years | 5,423 | 59.92 (8.77) | 2,040 | 60.12 (8.67) |
| 16-18 years | 3,498 | 64.15 (6.16) | 1,389 | 64.67 (6.16) |
| PRS | 6,734 | -0.003 (1.00) | 1,125 | 0.02 (1.00) |

DBP, diastolic blood pressure; mmHg, millimetres of mercury; PRS; polygenic risk score; SBP, systolic blood pressure; SD, standard deviation.

Table S3.4 Descriptive statistics of glucose and PRS by inclusion and exclusion criteria

|  | INCLUDED SAMPLE | | | | | EXCLUDED SAMPLE | | |
| --- | --- | --- | --- | --- | --- | --- | --- | --- |
| Outcome | **Participants (N)** | | | **Mean (mmol/l) (SD)** | | **Participants (N)** | **Mean (mmol/l) (SD)** | |
| Glucose |  | | |  | |  |  | |
| 7 years | | 4,291 | 4.18 (0.50) | | 1,190 | | | 4.18 (0.51) |
| 7-15 years | | 699 | 4.94 (0.34) | | 159 | | | 4.94 (0.36) |
| 15-18 years | | 3,454 | 5.12 (0.39) | | 1096 | | | 5.09 (0.42) |

mmol/l, millimoles per litre; PRS, polygenic risk scores; SD, standard deviation.

Table S3.5 Descriptive statistics of triglycerides, non-HDL-c and HDL-c and PRS by inclusion and exclusion criteria

|  | INCLUDED SAMPLE | | EXCLUDED SAMPLE | |
| --- | --- | --- | --- | --- |
| Outcome | **Participants (N)** | **Mean (mmol/l) (SD)** | **Participants (N)** | **Mean (mmol/l) (SD)** |
| Triglycerides |  |  |  |  |
| Birth | 2,909 | 0.57 (0.39) | 1861 | 0.56 (0.37) |
| 0-9 years | 4,145 | 1.04 (0.47) | 1143 | 1.04 (0.47) |
| 9-18 years | 5,085 | 0.95 (0.44) | 1531 | 0.94 (0.43) |
| PRS | 6,953 | 0.0003 (1.00) | 906 | 0.001 (0.96) |
| Non-HDL-c |  |  |  |  |
| Birth | 2,811 | 1.22 (0.53) | 1,809 | 1.21 (0.53) |
| 0-9 years | 4,164 | 2.87 (0.62) | 1,154 | 2.92 (0.64) |
| 9-18 years | 5,096 | 2.65 (0.66) | 1,542 | 2.64 (0.68) |
| PRS | 6,942 | -0.001 (1.00) | 917 | 0.01 (0.95) |
| HDL-c |  |  |  |  |
| Birth | 2,869 | 0.53 (0.24) | 1,831 | 0.53 (0.24) |
| 0-8 years | 3,972 | 1.52 (0.30) | 1056 | 1.52 (0.29) |
| 8-17 years | 5,163 | 1.34 (0.31) | 1429 | 1.32 (0.30) |
| PRS | 6,951 | -0.001 (1.00) | 908 | 0.01 (0.95) |

HDL-c, high-density lipoprotein; mmol/l, millimoles per litre; non-HDL-c, non-high-density lipoprotein; PRS, polygenic risk score; SD, standard deviation.

| Table S3.6 Descriptive statistics for CRP, PRS by inclusion and exclusion criteria | | | | |
| --- | --- | --- | --- | --- |
|  | **INCLUDED SAMPLE** | | **EXCLUDED SAMPLE** | |
| Outcome | **Participants (N)** | **Mean (mg/l) (SD)** | **Participants (N)** | **Mean (mg/l) (SD)** |
| 9 years | 4,018 | 0.57 (1.09) | 1,010 | 0.58 (1.03) |
| 9-18 years | 3,456 | 1.12 (2.02) | 1,093 | 1.20 (2.13) |
| PRS | 5,058 | 0.001 (1.00) | 2,801 | -0.001 (1.00) |

CRP, c-reactive protein; PRS, polygenic risk score; SD, standard deviation.

| Table S4.1 Model details for height trajectories included in the analysis | | | | | | |
| --- | --- | --- | --- | --- | --- | --- |
|  | No of contributing individuals | | Assessment of model fit | | | |
|  | Total number of observations | Number of individuals with 1 measure | Mean observed, cm (SD) | Mean predicted, (cm (SD)) | Mean difference (observed – predicted) | 95% level of agreement between observed and predicted |
| Total |  |  |  |  |  |  |
| 1 years | 9,330 | 5,379 | 81.19 (4.56) | 80.90 (3.96) | 0.29 | -3.06 to 3.63 |
| 5 years | 5,349 | 3,565 | 114.28 (5.99) | 114.42 (4.76) | -0.13 | -5.86 to 5.60 |
| 10 years | 6,554 | 5,375 | 143.14 (6.59) | 144.44 (6.43) | -1.29 | -4.12 to 1.54 |
| 17 years | 2,828 | 2,823 | 171.47 (9.31) | 174.07 (8.95) | -2.60 | -5.76 to 0.57 |
| cm, centimetres; SD, standard deviation. **Table S5** Characteristics at birth of the mothers of children included in models of glucose (risk factor with least individuals) \| **Inclusion sample^a^** **N= 4,291** \| \| **Exclusion sample^b^** **n=10,220** \| \| **P value for comparison^c^**; \| N \| N (%) \| N \| N (%) \|; **Sex of child** \| \| \| \| \| 0.796; Male \| 2,204 \| 51.44 \| 5,229 \| 51.20 \|; Female \| 2,081 \| 48.56 \| 4,984 \| 48.80 \|; **Maternal marital status** \| \| \| \| \| <0.0001; Never married \| 334 \| 9.20 \| 970 \| 14.92 \|; Widowed \| 8 \| 0.22 \| 20 \| 0.31 \|; Divorced \| 112 \| 3.08 \| 288 \| 4.43 \|; Separated \| 54 \| 1.49 \| 142 \| 2.18 \|; 1^st^ Marriage \| 2,827 \| 77.86 \| 4,568 \| 70.26 \|; Marriage 2 or 3 \| 296 \| 8.15 \| 514 \| 7.91 \|; **Household social class** \| \| \| \| \|; Professional \| 651 \| 19.56 \| 748 \| 12.31 \| <0.0001; Managerial & Technical \| 1,530 \| 45.97 \| 2,693 \| 44.32 \|; Non-manual \| 824 \| 24.76 \| 1,767 \| 29.08 \|; Manual \| 249 \| 7.48 \| 645 \| 10.62 \|; Part Skilled & Unskilled \| 74 \| 2.22 \| 223 \| 3.67 \|; **Maternal education** \| \| \| \| \|; Less than O level \| 786 \| 20.01 \| 2,896 \| 34.78 \| <0.0001; O level \| 1,350 \| 34.36 \| 2,889 \| 34.70 \|; A level \| 1,071 \| 27.26 \| 1,688 \| 20.27 \|; Degree or above \| 722 \| 18.38 \| 853 \| 10.25 \|; **Partners highest educational qualification** \| \| \| \| \|; Less than O level \| 990 \| 25.78 \| 3,082 \| 38.82 \| <0.0001; O level \| 835 \| 21.74 \| 1,666 \| 20.99 \|; A level \| 1,064 \| 27.71 \| 2,004 \| 25.24 \|; Degree or Above \| 951 \| 24.77 \| 1,187 \| 14.95 \|; **Maternal smoking during pregnancy** \| \| \| \| \|; Yes \| 698 \| 17.52 \| 2,547 \| 28.29 \| <0.0001; No \| 3,287 \| 82.48 \| 6,455 \| 71.71 \|; \| *Total (N)* \| *Mean (SD)* \| *Total* \| *Mean (SD)* \| P value; Child gestational age at birth \| 4,052 \| 39.50 (1.79) \| 9,740 \| 39.41 (1.93) \| 0.009 ^a^ Denominators for included participants in this table may be less than N included in full multilevel model due to missing data for these characteristics at baseline which were not required for our model (age, sex, at least one measure of risk factor were required for inclusion). Denominator for participants excluded may also vary due to missing data on the characteristics included in the table. ^b^ Exclusion sample includes participants who are singletons, alive at 1 year of age with no genetic or glucose data. ^c^ p value is for the difference in proportions for categorical variables from *χ*² test or difference in means for continuous variables from t tests between included and excluded participants | | | | | | |

| **Table S6.1** Cross-sectional analyses for the associations between Alzheimer’s disease PRS at *p*≤5x10^-8^ and birthweight | | | | | |
| --- | --- | --- | --- | --- | --- |
| **Outcome** | **N** | **β-coefficient**  **(95% CI)** | **β-coefficient/1 SD increase in PRS (95% CI)** | **p^a^** | **R^2^** |
| Birthweight (g) | 2,785 | 165.11 | 0.65 (-15.96,17.25) | 0.94 | 4.62x10^-5^ |

CI, confidence interval; g, grams; PRS, polygenic risk score; SD, standard deviation.

**^a^** P value is for the difference in the outcome per year where year is indicated for each 1 SD increase in PRS.

| **Table S6.2** The association between Alzheimer’s disease PRS at *p≤*5x10^-8^ and height | | | |
| --- | --- | --- | --- |
| Outcome | **Mean trajectory (95% CI)** | **Mean difference in risk factor**  **(95% CI) per 1 SD higher PRS** | **p^a^** |
| Height |  |  |  |
| *Age 1yr (cm)* | 75.69 (75.63, 75.75) | 0.01 (-0.05, 0.06) | 0.73 |
| Age 5 (cm) | 109.93 (109.82, 110.03) | 0.04 (-0.05, 0.14) | 0.38 |
| Age 10 (cm) | 140.44 (140.28, 140.60) | 0.08 (-0.06, 0.23) | 0.27 |
| Age 17 (cm) | 178.33 (178.12, 178.537) | 0.02 (-0.16, 0.19) | 0.86 |

CI, confidence interval; cm, centimetres; PRS, polygenic risk scores; SD, standard deviation.

**^a^** P value is for the difference in the outcome per year where year is indicated for each 1 SD increase in PRS.

**Table S6.3** The association between Alzheimer’s disease PRS at *p*≤5x10^-8^ and anthropometric risk factors

| Outcome | Mean trajectory (95% CI) | Mean difference in anthropometry (95% CI) per 1 SD higher PRS | p^c^ |
| --- | --- | --- | --- |
| Height-adjusted fat mass |  |  |  |
| *Age 9yr* | 2.01 ln(kg) (1.99, 2.04) **^a^** | 0.59% (-0.92, 2.11) **^b^** | 0.44 |
| Change 9-13yr | 0.15 ln(kg/yr) (0.15, 0.16) **^a^** | -0.15%/yr (-0.46, 0.16) **^b^** | 0.34 |
| Change 13-15yr | 0.11 ln(kg/yr) (0.10, 0.11) **^a^** | 0.07%/yr (-0.48, 0.63) **^b^** | 0.79 |
| Change 15-18yr | 0.06 ln(kg/yr) (0.05, 0.06) **^a^** | 0.17%/yr (-0.25, 0.58) **^b^** | 0.43 |
| Height-adjusted lean mass |  |  |  |
| *Age 9yr (kg)* | 20.94 kg (20.81, 21.07) | 0.04 kg (-0.03, 0.11) | 0.24 |
| Change 9-13yr (kg/yr) | 2.92 kg/yr (2.88, 2.97) | -0.01 kg/yr (-0.04, 0.02) | 0.69 |
| Change 13-15yr (kg/yr) | 2.35 kg/yr(2.25, 2.44) | 0.02 kg/yr (-0.05, 0.09) | 0.55 |
| Change 15-18yr (kg/yr) | 0.26 kg/yr (0.19, 0.32) | -0.01 kg/yr (-0.06, 0.04) | 0.71 |

Change/yr, change per year; CI, confidence interval; kg, kilograms; kg/yr, kilograms per year; yr, year; ln, natural logarithm; %/yr, percentage per year; PRS, polygenic risk score; SD, standard deviation.

**^a^** Height-adjusted fat mass was transformed using the natural log. All values are in log form.

**^b^** The difference in fat mass per 1 SD higher PRS is back transformed from the log scale for ease of interpretation and is interpreted as the percentage difference in the mean level or percentage difference in change per year.

**^C^** P value is for the difference in the outcome per year where year is indicated for each 1 SD increase in PRS.

### **Table S6.4** The association between Alzheimer’s disease PRS at *p*≤5x10^-8^, SBP, and DBP

| Outcome | Mean trajectory (95% CI) | Mean difference in risk factor (95% CI) per 1 SD higher PRS | p^a^ |
| --- | --- | --- | --- |
| SBP |  |  |  |
| *Age 7yr (mmHg)* | 98.53 (98.14, 98.92) | 0.07 (-0.16, 0.31) | 0.54 |
| Change 7-12yr (mmHg/yr) | 1.54 (1.46, 1.62) | -0.01 (-0.06, 0.05) | 0.83 |
| Change 12-16yr (mmHg/yr) | 3.35 (3.22, 3.48) | 0.03 (-0.06, 0.12) | 0.51 |
| Change 16-18yr (mmHg/yr) | -5.07 (-5.36, -4.79) | 0.01 (-0.20, 0.22) | 0.90 |
| DBP |  |  |  |
| *Age 7yr (mmHg)* | 57.14 (56.86, 57.42) | 0.19 (0.02, 0.37) | 0.03 |
| Change 7-12yr (mmHg/yr) | -0.002 (-0.06, 0.06) | -0.05 (-0.09, -0.003) | 0.03 |
| Change 12-16yr (mmHg/yr) | 2.12 (2.01, 2.23) | 0.07 (-0.01, 0.14) | 0.09 |
| Change 16-18yr (mmHg/yr) | -0.39 (-0.62, -0.17) | -0.04 (-0.21, 0.12) | 0.60 |

Change/yr, change per year; CI, confidence interval; DBP, diastolic blood pressure; mmHg, millimetres of mercury; mmHg/yr, millimetres of mercury per year; PRS, polygenic risk score; SBP, systolic blood pressure; SD, standard deviation; yr, year.

**^a^** P value is for the difference in the outcome per year where year is indicated for each 1 SD increase in PRS.

Table S6.5 The association between Alzheimer’s disease PRS at *p*≤5x10^-8^ and glucose

| Outcome | Mean trajectory (95% CI) | Mean difference in risk factor (95% CI) per 1 SD higher PRS | p^a^ |
| --- | --- | --- | --- |
| Glucose |  |  |  |
| *Age 7yr (mmol/l)* | 4.10 (4.08, 4.13) | 0.01 (-0.005, 0.03) | 0.18 |
| Change 7-15yr (mmol/l/yr) | 0.14 (0.13, 0.14) | 0.00001 (-0.003, 0.003) | 0.99 |
| Change 15-18yr (mmol/l/yr) | -0.10 (-0.11, -0.09) | -0.005 (-0.01, 0.003) | 0.25 |

Change/yr, change per year; CI, confidence interval; mmol/l, millimoles per litre; mmol/l/yr, millimoles per litre per year; PRS, polygenic risk scores; SD, standard deviation; yrs, years.

**^a^** P value is for the difference in the outcome per year where year is indicated for each 1 SD increase in PRS.

**Table S6.6** The association between Alzheimer’s disease PRS at *p*≤5x10^-8^, log triglycerides and cholesterol

| Outcome | Mean trajectory (95% CI) | Mean difference in lipids  (95% CI) per 1 SD higher PRS | p^c^ |
| --- | --- | --- | --- |
| Triglycerides |  |  |  |
| *Birth* | -0.66 ln(mmol/l) (-0.69, -0.64) **^a^** | -0.07% (-1.68, 1.53)**^b^** | 0.93 |
| Change 0-9yr | 0.08 ln(mmol/l/yr) (0.08, 0.09) **^a^** | 0.03%/yr (-0.20, 0.26)**^b^** | 0.80 |
| Change 9-18yr | -0.04 ln(mmol/l/yr) (-0.05, -0.04)**^a^** | -0.02%/yr (-0.20, 0.16)**^b^** | 0.84 |
| HDL-c |  |  |  |
| *Birth (mmol/l)* | 0.56 mmol/l/ (0.54, 0.57) | 0.003 mmol/l (-0.01, 0.01) | 0.48 |
| Change 0-7yr (mmol/l/yr) | 0.12 mmol/l/yr (0.11, 0.12) | -0.0002 mmol/l/yr (-0.002, 0.001) | 0.80 |
| Change 7-18yr (mmol/l/yr) | -0.02 mmol/l/yr (-0.02, -0.02) | -0.0001 mmol/l/yr (-0.001, 0.001) | 0.88 |
| Non-HDL-c |  |  |  |
| *Birth (mmol/l)* | 1.31 mmol/l (1.28, 1.34) | 0.01 mmol/l (-0.01, 0.02) | 0.58 |
| Change 0-9yr (mmol/l/yr) | 0.20 mmol/l/yr (0.20, 0.21) | -0.002 mmol/l /yr (-0.004, 0.001) | 0.29 |
| Change 9-18yr (mmol/l/yr) | -0.07 mmol/l/yr (-0.08, -0.07) | 0.002 mmol/l /yr (-0.001, 0.004) | 0.14 |

Change/yr, change per year; CI, confidence interval; HDL-c, high-density lipoprotein; ln, natural logarithm; %/yr, percentage per year; PRS, polygenic risk score; SD, standard deviation; yr, years.

**^a^** Triglycerides were transformed using the natural log. All values are in log form.

**^b^** The difference in triglyceride levels/1 SD increase in PRS is back transformed from the log scale for ease of interpretation and is interpreted as the percentage difference in the mean level or percentage change per year for a 1 SD increase in PRS.

**^c^** P value is for the difference in the outcome per year where year is indicated for each 1 SD increase in PRS.

**Table S6.7** The association between Alzheimer’s disease PRS at *p*≤5x10^-8^ and CRP

| Outcome | Mean trajectory  (95% CI) | Mean difference in CRP  (95% CI) per 1 SD higher PRS | p^c^ |
| --- | --- | --- | --- |
| CRP |  |  |  |
| *Age 9yr* | -1.25 ln(mg/l) (-1.31, -1.18)**^a^** | 0.07% (-3.65, 3.79)**^b^** | 0.97 |
| Change 9-18yr | 0.10 ln(mg/l/yr) (0.09, 0.11)**^a^** | 0.08%/yr (-0.51, 0.68)**^b^** | 0.78 |

Change/yr, change per year; CI, confidence interval; CRP, c-reactive protein; ln, natural logarithm; mg/l, milligrams per litre; mg/l/yr, milligrams per litre per year; PRS, polygenic risk scores; SD, standard deviation; yr, year.

**^a^** CRP was transformed using the natural log. All values are in log form.

**^b^** The difference in CRP/1 SD increase in PRS is back transformed from the log scale for ease of interpretation and is interpreted as the percentage difference in mean CRP/1 SD increase in PRS.

**^c^** P value is for the difference in the outcome per year where year is indicated for each 1 SD increase in PRS.

| **Table S6.8** Cross-sectional analyses for the association between Alzheimer’s disease PRS at *p*≤5x10^-8^, IL-6, and insulin | | | | | |
| --- | --- | --- | --- | --- | --- |
| **Outcome** | **N** | **β-coefficient**  **(95% CI)** | **β-coefficient/1 SD increase in PRS**  **(95% CI)** | **p^c^** | **R^2^** |
| Age 9yr (IL-6) | 4,051 | 0.02 ln(pg/ml) (0.01, 0.02)**^a^** | -1.95% (-4.52, 0.70)**^b^** | 0.15 | 2.41x10^-3^ |
| Age 15yr (Insulin) | 2,774 | 0.01 ln(mu/l) (0.002, 0.01)**^a^** | 1.15% (-0.61, 2.94)**^b^** | 0.20 | 3.90x10^-3^ |

CI, confidence interval; ln, natural logarithm; mu/l, milliunits per litre; pg/ml, picograms per millilitre; PRS, polygenic risk scores; SD, standard deviation.

**^a^** Insulin and IL-6 were transformed using the natural log. All values are in log form.

**^b^** The difference in outcome/1 SD increase in PRS is back transformed from the log scale for ease of interpretation and is interpreted as the percentage difference in the mean level or percentage difference in change per year.

**^c^** P value is for the difference in the outcome for each 1 SD increase in PRS.

| **Table S6.9** Cross-sectional analyses for the association between Alzheimer’s disease PRS at *p*≤5x10^-8^ and MVPA | | | | | |
| --- | --- | --- | --- | --- | --- |
| **Outcome** | **N** | **β-coefficient**  **(95% CI)** | **β-coefficient/1 SD increase in PRS**  **(95% CI)** | **p^c^** | **R^2^** |
| **MVPA** |  |  |  |  |  |
| Age 11yr | 4,389 | -0.01 ln(mins/day) (-0.02, -0.004)**^a^** | -0.25% (-2.36, 1.91)**^b^** | 0.82 | 4.34 x10^-4^ |
| Age 12yr | 2,435 | -0.001 ln(mins/day) (-0.001, 0.00002)**^a^** | -3.65% (-8.09, 1.01)**^b^** | 0.12 | 1.07x10^-3^ |
| Age 15yr | 1,805 | -0.01 ln(mins/day) (-0.02, -0.01)**^a^** | -0.23% (-4.59, 4.34)**^b^** | 0.92 | 1.27x10^-3^ |

CI, confidence interval; ln, natural logarithm; mins/day, minutes per day; MVPA, moderate to vigorous physical activity; PRS, polygenic risk score; SD, standard deviation.

**^a^** MVPA was transformed using the natural log. All values are in log form.

**^b^** The difference in MVPA per 1 SD higher PRS is back transformed from the log scale for ease of interpretation and is interpreted as the percentage difference in the mean level or percentage change per year for a 1 SD increase in PRS.

**^c^** P value is for the difference in the outcome for each 1 SD increase in PRS.

| **Table S7.1** Cross-sectional analyses for the associations between Alzheimer’s disease PRS at *p*≤5x10^-2^ and birthweight | | | | |
| --- | --- | --- | --- | --- |
| **Outcome** | **N** | **Mean difference in birthweight per 1 SD increase in PRS (95% CI)** | **p^a^** | **R^2^** |
| Birthweight (g) | 2,785 | 9.59 (-7.28, 26.45) | 0.27 | 3.48x10^-4^ |

CI, confidence interval; g, grams; PRS, polygenic risk score; SD, standard deviation.

**^a^** P value is for the difference in the outcome for each 1 SD increase in PRS.

Table S7.2 The association between Alzheimer’s disease PRS at *p*≤5x10^-2^ and height

| Outcome | Mean difference in height (95% CI) per 1 SD higher PRS | p^a^ |
| --- | --- | --- |
| Height |  |  |
| *Age 1yr (cm)* | -0.03 (-0.08, 0.03) | 0.34 |
| Age 5yr (cm) | -0.10 (-0.19, -0.003) | 0.04 |
| Age 10yr (cm) | -0.15 (-0.30, -0.01) | 0.04 |
| Age 17yr (cm) | -0.04 (-0.21, 0.14) | 0.67 |

CI, confidence interval; cm, centimetres; PRS, polygenic risk scores; SD, standard deviation.

**^a^** P value is for the difference in the outcome per year where year is indicated for each 1 SD increase in PRS.

**Table S7.3** The association between Alzheimer’s disease PRS at *p*≤5x10^-2^ and anthropometric risk factors

| **Outcome** | **Mean difference in anthropometry (95% CI) per 1 SD higher PRS** | **p^b^** |
| --- | --- | --- |
| **Height-adjusted fat mass** |  |  |
| *Age 9yr* | -0.68% (-2.16, 0.80)**^a^** | 0.37 |
| Change 9-13yr | 0.02%/yr (-0.29, 0.33)**^a^** | 0.90 |
| Change 13-15yr | 0.26%/yr (-0.29, 0.81)**^a^** | 0.36 |
| Change 15-18yr | 0.01%/yr (-0.41, 0.42)**^a^** | 0.98 |
| **Height-adjusted lean mass** |  |  |
| *Age 9yr (kg)* | -0.07 kg (-0.14, 0.004) | 0.06 |
| Change 9-13yr (kg/yr) | 0.01kg/yr (-0.02, 0.04) | 0.61 |
| Change 13-15yr (kg/yr) | 0.08kg/yr (0.02, 0.15) | 0.02 |
| Change 15-18yr (kg/yr) | -0.06 kg/yr (-0.11, -0.02) | 0.01 |

Change/yr, change per year; CI, confidence interval; kg, kilograms; kg/yr, kilograms per year; %/yr,

**^a^** The difference in fat mass per 1 SD higher PRS is back transformed from the log scale for ease of interpretation and is interpreted as the percentage difference in the mean level or percentage change per year for a 1 SD increase in PRS.

**^b^** P value is for the difference in the outcome per year where year is indicated for each 1 SD increase in PRS.

**Table S7.4** The association between Alzheimer’s disease PRS at *p*≤5x10^-2^, SBP, and DBP

| **Outcome** | **Mean difference in blood pressure (95% CI) per 1 SD higher PRS** | **p^a^** |
| --- | --- | --- |
| **SBP** |  |  |
| *Age 7yr (mmHg)* | -0.09 (-0.32, 0.15) | 0.48 |
| Change 7-12yr (mmHg/yr) | 0.01 (-0.05, 0.07) | 0.71 |
| Change 12-16yr (mmHg/yr) | 0.05 (-0.04, 0.14) | 0.31 |
| Change 16-18yr (mmHg/yr) | -0.14 (-0.35, 0.07) | 0.18 |
| **DBP** |  |  |
| *Age 7yr (mmHg)* | 0.14 (-0.03, 0.31) | 0.11 |
| Change 7-12yr (mmHg/yr) | -0.02 (-0.06, 0.02) | 0.40 |
| Change 12-16yr (mmHg/yr) | 0.01 (-0.07, 0.09) | 0.82 |
| Change 16-18yr (mmHg/yr) | 0.002 (-0.16, 0.17) | 0.98 |

Change/yr, change per year; CI, confidence interval; DBP, diastolic blood pressure; mmHg, millimetres of mercury; mmHg/yr, millimetres of mercury per year; PRS, polygenic risk score; SBP, systolic blood pressure; SD, standard deviation; yr, year.

**^a^** P value is for the difference in the outcome per year where year is indicated for each 1 SD increase in PRS.

Table S7.5 The association between Alzheimer’s disease PRS at *p*≤5x10^-2^ and glucose

| Outcome | Mean difference in glucose (95% CI) per 1 SD higher PRS | p^a^ |
| --- | --- | --- |
| Glucose |  |  |
| *Age 7yr (mmol/l)* | 0.01 (-0.002, 0.03) | 0.08 |
| Change 7-15yr (mmol/l/yr) | 0.0002 (-0.003, 0.003) | 0.87 |
| Change 15-18yr (mmol/l/yr) | -0.003 (-0.01, 0.01) | 0.54 |

Change/yr, change per year; CI, confidence interval; mmol/l, millimole per litre; mmol/l/year, millimoles per litre per year; yr, year; SD, standard deviation.

**^a^** P value is for the difference in the outcome per year where year is indicated for each 1 SD increase in PRS.

**Table S7.6** The association between Alzheimer’s disease PRS at *p*≤5x10^-2^, log triglycerides and cholesterol

| Outcome | Mean difference in lipids (95% CI) per 1 SD higher PRS | p^b^ |
| --- | --- | --- |
| Triglycerides |  |  |
| *Birth* | 2.07% (0.42, 3.72)**^a^** | 0.01 |
| Change 0-9yr | -0.23%/yr (-0.46, -0.003**) ^a^** | 0.05 |
| Change 9-18yr | 0.07%/yr (-0.11, 0.25) **^a^** | 0.43 |
| HDL-c |  |  |
| *Birth (mmol/l)* | 0.003 mmol/l (-0.01, 0.01) | 0.50 |
| Change 0-7yr (mmol/l/yr) | 0.0002 mmol/l/yr (-0.001, 0.002) | 0.74 |
| Change 7-18yr (mmol/l/yr) | -0.001 mmol/l/yr (-0.002, 0.0004) | 0.21 |
| Non-HDL-c |  |  |
| *Birth (mmol/l)* | 0.01mmol/l (-0.004, 0.03) | 0.12 |
| Change 0-9yr (mmol/l/yr) | -0.002 mmol/l/yr (-0.005, 0.001) | 0.21 |
| Change 9-18yr (mmol/l/yr) | 0.001 mmol/l/yr (-0.002, 0.003) | 0.68 |

Change/yr, change per year; CI, confidence interval; HDL-c, high-density lipoprotein; %/yr, percentage per year; PRS, polygenic risk score; SD, standard deviation; yr, years.

**^a^** The difference in triglyceride levels/1 SD increase in PRS is back transformed from the log scale for ease of interpretation and is interpreted as the percentage difference in the mean level or percentage change per year for a 1 SD increase in PRS.

**^b^** P value is for the difference in the outcome per year where year is indicated for each 1 SD increase in PRS.

**Table S7.7** The association between Alzheimer’s disease PRS at *p*≤5x10^-2^ and CRP

| Outcome | Mean difference in CRP (95% CI) per 1 SD higher PRS | p^b^ |
| --- | --- | --- |
| CRP |  |  |
| *Age 9yr* | -3.82% (-7.36, -0.29)**^a^** | 0.04 |
| Change 9-18yr | 0.21%/yr (-0.38, 0.80)**^a^** | 0.48 |

Change/yr, change per year; CI, confidence interval; CRP, c-reactive protein; mg/l, milligrams per litre; mg/l/yr, milligrams per litre per year; PRS, polygenic risk score; SD, standard deviation; yr, year(s).

**^a^** The difference in CRP/1 SD increase in PRS is back transformed from the log scale for ease of interpretation and is interpreted as the percentage difference in mean CRP/1 SD increase in PRS.

**^b^** P value is for the difference in the outcome per year where year is indicated for each 1 SD increase in PRS.

| **Table S7.8** Association between Alzheimer’s disease PRS at *p*≤5x10^-2^, IL-6 and insulin | | | | |
| --- | --- | --- | --- | --- |
| **Outcome** | **N** | **Mean difference in outcome (95% CI) per 1 SD higher PRS** | **p^b^** | **R^2^** |
| Age 9yr (IL-6) | 4,051 | 0.03% (-2.58, 2.71)**^a^** | 0.98 | 1.91x10^-3^ |
| Age 15yr (Insulin) | 2,774 | 2.30% (0.54, 4.08)**^a^** | 0.01 | 5.63x10^-3^ |

CI, confidence interval; IL-6, interleukin-6; pg/ml, picograms per millilitre; PRS, polygenic risk score; SD, standard deviation.

**^a^** The difference in outcome/1 SD increase in PRS is back transformed from the log scale for ease of interpretation and is interpreted as the percentage difference in mean outcome/1 SD increase in PRS.

**^b^** P value is for the difference in the outcome for each 1 SD increase in PRS.

| **Table S7.9** Cross-sectional analyses for the association between the PRS for Alzheimer’s disease at *p*≤5x10^-2^ and MVPA | | | | |
| --- | --- | --- | --- | --- |
| **Outcome** | **N** | **Mean difference in outcome (95% CI) per 1 SD higher PRS** | **p^b^** | **R^2^** |
| **MVPA** |  |  |  |  |
| Age 12yrs | 4,389 | 1.27% (-0.87, 3.45)**^a^** | 0.25 | 6.99 x10^-4^ |
| Age 14yrs | 3,281 | 1.15% (-3.56, 6.09) **^a^** | 0.64 | 2.04x10^-4^ |
| Age 15yrs | 1,818 | 3.25% (-1.18, 7.87)**^a^** | 0.15 | 2.32x10^-3^ |

CI, confidence interval; mins/day, minutes per day; MVPA, moderate to vigorous physical activity; PRS, polygenic risk score; SD, standard deviation.

**^a^** The difference in MVPA per 1 SD higher PRS is back transformed from the log scale for ease of interpretation and is interpreted as the percentage difference in the mean level or percentage change per year for a 1 SD increase in PRS.

**^b^** P value is for the difference in the outcome for each 1 SD increase in PRS.

| **Table S8.1** Cross-sectional analyses for the associations between the PRS for Alzheimer’s disease at *p*≤5x10^-1^ and birthweight | | | |  |  |
| --- | --- | --- | --- | --- | --- |
| **Outcome** | **N** | **Mean difference in outcome (95% CI) per 1 SD higher PRS** | **p^a^** | | **R^2^** |
| Birthweight (g) | 2,785 | 0.77 (-15.89, 17.42) | 0.93 | | 4.68x10^-5^ |

CI, confidence interval; g, grams; PRS, polygenic risk score; SD, standard deviation.

**^a^** P value is for the difference in the outcome for each 1 SD increase in PRS.

| **Table S8.2** The association between Alzheimer’s disease PRS at *p*≤5x10^-1^ and height | | |
| --- | --- | --- |
| Outcome | **Mean difference in height (95% CI) per 1 SD higher PRS** | **p^a^** |
| Height |  |  |
| *Age 1yr (cm)* | -0.03 (-0.08, 0.03) | 0.30 |
| Age 5 (cm) | -0.10 (-0.20, 0.-0.01) | 0.03 |
| Age 10 (cm) | -0.15 (-0.29, -0.004) | 0.04 |
| Age 17 (cm) | -0.14 (-0.31, 0.04) | 0.13 |

CI, confidence interval; cm, centimetre; PRS, polygenic risk scores; SD, standard deviation.

**^a^** P value is for the difference in the outcome per year where year is indicated for each 1 SD increase in PRS.

| Table S8.3 The association between Alzheimer’s disease PRS at *p*≤5x10^-1^ and anthropometric factors | | |
| --- | --- | --- |
| **Outcome** | **Mean difference in anthropometry (95% CI) per 1 SD higher PRS** | **p^b^** |
| **Height-adjusted fat mass** |  |  |
| *Age 9yr (%)* | -0.78% (-2.25, 0.70) **^a^** | 0.31 |
| Change 9-13yr (%/yr) | 0.05%/yr (-0.26, 0.36)**^a^** | 0.75 |
| Change 13-15yr (%/yr) | 0.07%/yr (-0.48, 0.62) **^a^** | 0.81 |
| Change 15-18yr (%/yr) | 0.08%/yr (-0.34, 0.49)**^a^** | 0.72 |
| **Height-adjusted lean mass** |  |  |
| *Age 9yr (kg)* | -0.08kg (-0.15, -0.01) | 0.02 |
| Change 9-13yr (kg/yr) | 0.01kg/yr (-0.02, 0.04) | 0.43 |
| Change 13-15yr (kg/yr) | 0.05kg/yr (-0.02, 0.11) | 0.16 |
| Change 15-18 (kg/yr) | -0.06kg/yr (-0.11, -0.02) | 0.01 |

Change/yr, change per year; CI, confidence interval; kg, kilograms; kg/yr, kilograms per year; %/yr, percentage per year; PRS, polygenic risk score; standard deviation; yr, year(s).

**^a^** The difference in fat mass per 1 SD higher PRS is back transformed from the log scale for ease of interpretation and is interpreted as the percentage difference in the mean level or percentage change per year for a 1 SD increase in PRS.

**^b^** P value is for the difference in the outcome per year where year is indicated for each 1 SD increase in PRS.

### **Table S8.4** The association between Alzheimer’s disease PRS at *p*≤5x10^-1^ and blood pressure

| **Outcome** | **Mean difference in blood pressure (95% CI) per 1 SD higher PRS** | **p^a^** | |
| --- | --- | --- | --- |
| **SBP** |  | |  |
| *Age 7yr (mmHg)* | -0.05 (-0.29, 0.19) | | 0.68 |
| Change 7-12yr (mmHg/yr) | 0.01 (-0.05, 0.06) | | 0.80 |
| Change12-16yr (mmHg/yr) | 0.06 (-0.03, 0.15) | | 0.18 |
| Change 16-18yr (mmHg/yr) | -0.15 (-0.35, 0.06) | | 0.17 |
| **DBP** |  | |  |
| *Age 7yr (mmHg)* | 0.13 (-0.04, 0.30) | | 0.14 |
| Change 7-12yr (mmHg/yr) | -0.01 (-0.06, 0.03) | | 0.50 |
| Change 12-16yr (mmHg/yr) | 0.01 (-0.07, 0.09) | | 0.82 |
| Change 16-18yr (mmHg/yr) | -0.01 (-0.18, 0.15) | | 0.88 |

Change/yr, change per year; CI, confidence interval; DBP, diastolic blood pressure; mmHg, millimetres of mercury; mmHg/yr, millimetres of mercury per year; PRS, polygenic risk score; SBP, systolic blood pressure; standard deviation; SD; yr, year (s).

**^a^** P value is for the difference in the outcome per year where year is indicated for each 1 SD increase in PRS.

Table S8.5 The association between Alzheimer’s disease PRS at *p*≤5x10^-1^ and glucose

| **Outcome** | **Mean difference in glucose (95% CI) per 1 SD higher PRS** | **p^a^** |
| --- | --- | --- |
| **Glucose** |  |  |
| *Age 7yr (mmol/l)* | 0.01 (-0.01, 0.03) | 0.19 |
| Change 7-15yr (mmol/l/yr) | -0.001 (-0.004, 0.002) | 0.61 |
| Change 15-18yr (mmol/l/yr) | 0.0001 (-0.01, 0.01) | 0.99 |

Change/yr, change per year; CI, confidence interval; mmol/l, millimole per litre; mmol/l/year, millimoles per litre per year; SD, standard deviation; yr, year(s).

**^a^** P value is for the difference in the outcome per year where year is indicated for each 1 SD increase in PRS.

Table S8.6 The association between Alzheimer’s disease PRS at *p*≤5x10^-1^, log triglycerides and cholesterol

| **Outcome** | **Mean difference in lipids (95% CI) per 1 SD higher PRS** | **p^b^** |
| --- | --- | --- |
| **Triglycerides** |  |  |
| *Birth* | 1.75% (-0.11, 3.40)**^a^** | 0.03 |
| Change 0-9yr | -0.21%/yr (-0.44, 0.02)**^a^** | 0.07 |
| Change 9-18yr | 0.13%/yr (-0.05, 0.31)**^a^** | 0.17 |
| **Non-HDL-c** |  |  |
| *Birth (mmol/l)* | 0.01 mmol/l (-0.01, 0.03) | 0.21 |
| Change 0-9yr (mmol/l/yr) | -0.002 mmol/l/yr (-0.005, 0.001) | 0.28 |
| Change 9-18yr (mmol/l/yr) | 0.001 mmol/l/yr (-0.002, 0.003) | 0.58 |
| **HDL-c** |  |  |
| *Birth (mmol/l)* | 0.004 mmol/l (-0.005, 0.01) | 0.39 |
| Change 0-8yr (mmol/l/yr) | 0.0002 mmol/l/yr (-0.001, 0.002) | 0.81 |
| Change 8-18yr (mmol/l/yr) | -0.0004 mmol/l/yr(-0.001, 0.001) | 0.47 |
| Change/yr, change per year; CI, confidence interval; HDL-c, high-density lipoprotein; mmol/l, millimole per litre; mmol/l, millimole per litre per year; %/yr, percentage per year; PRS, polygenic risk score; standard deviation; yr, year(s).  **^a^** The difference in triglycerides per 1 SD higher PRS is back transformed from the log scale for ease of interpretation and is interpreted as the percentage difference in the mean level or percentage change per year for a 1 SD increase in PRS.  **^b^** P value is for the difference in the outcome per year where year is indicated for each 1 SD increase in PRS. | | |

### **Table S8.7** The association between Alzheimer’s disease PRS at *p*≤5x10^-1^ and log CRP

| **Outcome** | **Mean difference in CRP (95% CI) per 1 SD higher PRS** | **p^b^** |
| --- | --- | --- |
| **CRP** |  |  |
| *Age 9yr (%)* | -3.24% (-6.82, 0.35)**^a^** | 0.08 |
| Change 9-18 (%/yr) | 0.04%/yr (-0.55, 0.63)**^a^** | 0.74 |

Change/yr, change per year; CI, confidence interval; CRP, C-reactive protein; %/yr, percentage per year; PRS, polygenic risk score; SD, standard deviation; yr. year(s).

**^a^** The difference CRP/1 SD increase in PRS is back transformed from the log scale for ease of interpretation and is interpreted as the percentage difference in mean CRP/1 SD increase in PRS.

**^b^** P value is for the difference in the outcome per year where year is indicated for each 1 SD increase in PRS.

**Table S8.8** Association between Alzheimer’s disease PRS at *p*≤5x10^-1^, IL-6 and insulin

| **Outcome** | **N** | **Mean difference in IL-6 (95% CI) per 1 SD higher PRS** | **p ^b^** | **R^2^** |
| --- | --- | --- | --- | --- |
| Age 9yr | 4,051 | 0.05% (-2.57, 2.73)**^a^** | 0.97 | 1.91x10^-3^ |
| Age 15yr | 2,774 | 1.58% (-0.16, 3.35) **^a^** | 0.08 | 4.44x10^-3^ |
| CI, confidence interval; IL-6, interleukin-6; PRS, polygenic risk score; SD, standard deviation; yr, year(s). **^a^** The difference in outcome/1 SD increase in PRS is back transformed from the log scale for ease of interpretation and is interpreted as the percentage difference in mean outcome/1 SD increase in PRS.  **^b^** P value is for the difference in the outcome for each 1 SD increase in PRS. | | | | |

| **Table S8.9:** Cross-sectional analyses for the association between the PRS for Alzheimer’s disease at *p*≤5x10^-1^ and MVPA | | | | |
| --- | --- | --- | --- | --- |
| **Outcome** | **N** | **Mean difference in MVPA per 1 SD higher PRS (95% CI)** | **p^b^** | **R^2^** |
| **MVPA** |  |  |  |  |
| Age 12yr | 4,389 | 0.70% (-1.41,2.86)**^a^** | 0.52 | 5.10x10^-4^ |
| Age 14yr | 2,435 | -2.00% (-6.57, 2.78)**^a^** | 0.41 | 3.92x10^-4^ |
| Age 15yr | 1,805 | 3.99% (-0.59, 8.79)**^a^** | 0.09 | 2.76x10^-3^ |
| CI, confidence interval; ln, natural logarithm; mins/day, minutes per day; MVPA, moderate to vigorous physical activity; PRS, polygenic risk score; SD, standard deviation; yrs, years.  **^a^** The difference in MVPA per 1 SD higher PRS is back transformed from the log scale for ease of interpretation and is interpreted as the percentage difference in the mean level or percentage change per year for a 1 SD increase in PRS.  **^b^** P value is for the difference in the outcome for each 1 SD increase in PRS. | | | | |

| **Table S9.1** Cross-sectional analyses for the associations between Alzheimer’s disease PRS (including the ApoE region) at *p*≤5x10^-8^ and birthweight | | | | |
| --- | --- | --- | --- | --- |
| **Outcome** | **N** | **Mean difference in birthweight per 1 SD increase in PRS (95% CI)** | **p^a^** | **R^2^** |
| Birthweight (g) | 2,785 | 11.06 (-5.95, 28.07) | 0.20 | 4.41x10^-4^ |

ApoE, apolipoprotein E; CI, confidence interval; g, grams; PRS, polygenic risk score; SD, standard deviation.

**^a^** P value is for the difference in the outcome for each 1 SD increase in PRS.

| **Table S9.2** The association between Alzheimer’s disease PRS (including the ApoE region) at *p*≤5x10^-8^ and height | | | |
| --- | --- | --- | --- |
| Outcome | **Mean trajectory (95% CI)** | **Mean difference in height (95% CI) per 1 SD higher PRS** | **p^a^** |
| Height |  |  |  |
| *Age 1yr (cm)* | 75.69 (75.63, 75.75) | 0.04 (-0.01, 0.10) | 0.12 |
| Age 5 (cm) | 109.93 (109.82, 110.03) | 0.07 (-0.03, 0.16) | 0.18 |
| Age 10 (cm) | 140.44 (140.28, 140.60) | 0.07 (-0.07, 0.22) | 0.31 |
| Age 17 (cm) | 178.33 (178.12, 178.537) | 0.09 (-0.08, 0.27) | 0.31 |

ApoE, apolipoprotein E; cm, centimetres; CI, confidence interval; PRS, polygenic risk scores; SD, standard deviation.

**^a^** P value is for the difference in the outcome per year where year is indicated for each 1 SD increase in PRS.

### Table S9.3 The association between Alzheimer’s disease PRS (including the ApoE region) at *p*≤5x10^-8^ and anthropometric risk factors

| **Outcome** | **Mean difference in anthropometry (95% CI) per 1 SD higher PRS** | **p ^b^** |
| --- | --- | --- |
| **Height-adjusted fat mass** |  |  |
| *Age 9yr (%)* | 0.06% (-1.42, 1.56)**^a^** | 0.94 |
| Change 9-13yr (%/yr) | -0.07%/yr (-0.38, 0.24)**^a^** | 0.66 |
| Change 13-15yr (%/yr) | 0.38% (-0.17, 0.94) **^a^** | 0.18 |
| Change 15-18yr (%/yr) | -0.13% (-0.54, 0.29) **^a^** | 0.54 |
| **Height-adjusted lean mass** |  |  |
| *Age 9yr (kg)* | 0.01kg (-0.06, 0.08) | 0.77 |
| Change 9-13yr (kg/yr) | -0.01 kg/yr (-0.04, 0.02) | 0.59 |
| Change 13-15yr (kg/yr) | -0.01 kg/yr (-0.08, 0.05) | 0.71 |
| Change 15-18yr (kg/yr) | -0.0002 kg/yr (-0.05, 0.05) | 0.99 |

ApoE, apolipoprotein E; CI, confidence interval; kg, kilograms; kg/yr, kilograms per year; %/yr, percentage per year; PRS, polygenic risk score; SD, standard deviation yr, year(s).

**^a^** The difference in fat mass per 1 SD higher PRS is back transformed from the log scale for ease of interpretation and is interpreted as the percentage difference in the mean level or percentage change per year for a 1 SD increase in PRS.

**^b^** P value is for the difference in the outcome per year where year is indicated for each 1 SD increase in PRS.

### **Table S9.4** The association between Alzheimer’s disease PRS (including the ApoE region) at *p*≤5x10^-8^ and blood pressure

| **Outcome** | **Mean difference in blood pressure (95% CI) per 1 SD higher PRS** | **p^a^** |
| --- | --- | --- |
| **SBP** |  |  |
| *Age 7yr (mmHg)* | -0.04 (-0.28, 0.19) | 0.73 |
| Change 7-12 (mmHg/yr) | 0.001 (-0.06, 0.06) | 0.97 |
| Change12-16 (mmHg/yr) | 0.07 (-0.02, 0.17) | 0.11 |
| Change 16-18 (mmHg/yr) | -0.14 (-0.35, 0.07) | 0.18 |
| **DBP** |  |  |
| *Age 7yr (mmHg)* | -0.21 (-0.38, -0.04)) | 0.02 |
| Change 7-12 (mmHg/yr) | 0.03 (-0.02, 0.07) | 0.22 |
| Change12-16 (mmHg/yr) | 0.05 (-0.02, 0.13) | 0.18 |
| Change 16-18 (mmHg/yr) | -0.12 (-0.29, 0.04) | 0.14 |

ApoE, apolipoprotein E; CI, confidence interval; mmHg, millimetres of mercury; mmHg/yr, millimetres of mercury per year; PRS, polygenic risk score; SD, standard deviation; yr, year.

**^a^** P value is for the difference in the outcome per year where year is indicated for each 1 SD increase in PRS.

Table S9.5 The association between Alzheimer’s disease PRS (including the ApoE region) and glucose at *p*≤5x10^-8^

| **Outcome** | **Mean difference in glucose (95% CI) per 1 SD higher PRS** | | **p^a^** |
| --- | --- | --- | --- |
| **Glucose (mmol/L)** |  |  | |
| *Age 7yr (mmol/l)* | -0.01 (-0.03, 0.005) | | 0.17 |
| Change 7-15yr (mmol/l/yr) | 0.001 (-0.002, 0.004) | | 0.40 |
| Change 15-18yr (mmol/l/yr) | -0.00002 (-0.01, 0.01) | | 0.99 |

ApoE, apolipoprotein E; CI, confidence interval; mmol/l, millimole per litre; mmol/l/year, millimoles per litre per year; SD, standard deviation; yr, year(s).

**^a^** P value is for the difference in the outcome per year where year is indicated for each 1 SD increase in PRS.

**Table S9.6** The association between Alzheimer’s disease PRS, cholesterol, and log triglycerides and PRS (including the ApoE region) at *p*≤5x10^-8^

| **Outcome** | **Mean difference in lipids (95% CI) per 1 SD higher PRS** | | **p^b^** |
| --- | --- | --- | --- |
| **Triglycerides** |  |  | |
| *Birth (%)* | 1.46% (-0.16, 3.10)**^a^** | | 0.08 |
| Change 0-9yr (%/yr) | 0.08%/yr (-0.15, 0.31)**^a^** | | 0.49 |
| Change 9-18yr (%/yr) | -0.12%/yr (-0.30, 0.06) **^a^** | | 0.20 |
| **Non-HDL-c** |  | |  |
| *Birth (mmol/l)* | 0.03 mmol/l (0.01, 0.04) | | 0.01 |
| Change 0-9yr (mmol/l/yr) | 0.01 mmol/l/yr (0.01, 0.02) | | <0.001 |
| Change 9-18yr (mmol/l/yr) | -0.003 mmol/l/yr (-0.01, -0.0005) | | 0.02 |
| **HDL-c** |  | |  |
| *Birth (mmol/l)* | -0.01mmol/l (-0.02, -0.005) | | 0.002 |
| Change 0-7yr (mmol/l/yr) | -0.002 mmol/l/yr (-0.003, -0.0004) | | 0.01 |
| Change 7-18yr (mmol/l/yr) | 0.001 mmol/l/yr (0.0002, 0.002) | | 0.02 |

ApoE, apolipoprotein E; CI, confidence interval; Change/yr, change per year.; HDL-c, high-density lipoprotein; mmol/l, millimole per litre; mmol/l, millimole per litre per year; %/yr, percentage per year; PRS, polygenic risk score; SD, standard deviation; yr, year(s).

**^a^** The difference in triglycerides per 1 SD higher PRS is back transformed from the log scale for ease of interpretation and is interpreted as the percentage difference in the mean level or percentage change per year for a 1 SD increase in PRS.

**^b^** P value is for the difference in the outcome per year where year is indicated for each 1 SD increase in PRS.

### **Table S9.7** The association between Alzheimer’s disease PRS (including the ApoE region) at *p*≤5x10^-8^ and CRP

| **Outcome** | **Mean difference in CRP (95% CI) per 1 SD higher PRS** | | **p^b^** |
| --- | --- | --- | --- |
| **CRP** |  |  | |
| *Age 9yr (CRP, %)* | -8.60% (-11.86, -5.21)**^a^** | 0.000001 | |
| Change 9-18 (CRP, %/yr) | 0.25%/yr (-0.33, 0.84) **^a^** | 0.39 | |

ApoE, apolipoprotein E; CI, confidence interval; CRP, c-reactive protein; %/yr, percentage per year; PRS, polygenic risk score; SD, standard deviation.

**^a^** CRP were transformed using the natural log. The difference in CRP per 1 SD higher PRS is back transformed from the log scale for ease of interpretation and is interpreted as the percentage difference in the mean level or percentage change per year for a 1 SD increase in PRS.

**^b^** P value is for the difference in the outcome per year where year is indicated for each 1 SD increase in PRS.

### Table S9.8 Association between Alzheimer’s disease PRS (including the ApoE region) at *p*≤5x10^-8^ and IL-6 at age 9

| **Outcome** | **N** | **Mean difference in IL-6 (95% CI) per 1 SD higher PRS** | **p^b^** | **R^2^** |
| --- | --- | --- | --- | --- |
| Age 9yr | 4,051 | 1.34% (-1.29, 4.03)**^a^** | 0.32 | 2.14x10^-3^ |
| Age 15yr | 2,774 | 0.74% (-1.00, 2.50)**^a^** | 0.41 | 3.58x10^-3^ |

ApoE, apolipoprotein E; CI, confidence interval; IL-6, interleukin-6; PRS, polygenic risk score; SD, standard deviation; yr, year(s).

**^a^** The difference in outcome/1 SD increase in PRS is back transformed from the log scale for ease of interpretation and is interpreted as the percentage difference in mean outcome/1 SD increase in PRS **^b^** P value is for the difference in the outcome for each 1 SD increase in PRS.

**Table 9.9** Cross-sectional analyses for the association between the PRS for Alzheimer’s disease (including the ApoE region) at *p*≤5x10^-8^ and MVPA

| **Outcome** | **N** | **Mean difference in MVPA (95% CI) per 1 SD higher PRS** | **p^b^** | **R^2^** |
| --- | --- | --- | --- | --- |
| **MVPA** |  |  |  |  |
| Age 11yr | 4,389 | 0.10% (-2.00, 2.24)**^a^** | 0.52 | 4.25x10^-4^ |
| Age 12yr | 2,435 | -0.98% (-5.50, 3.76)**^a^** | 0.67 | 1.83x10^-4^ |
| Age 15yr | 1,805 | -2.62% (-6.78, 1.73)**^a^** | 0.23 | 2.00x10^-3^ |

ApoE, apolipoprotein E; CI, confidence interval; ln, natural logarithm; mins/day, minutes per day; MVPA, moderate to vigorous physical activity; PRS, polygenic risk score; SD, standard deviation; yrs, years.

**^a^** The difference in MVPA per 1 SD higher PRS is back transformed from the log scale for ease of interpretation and is interpreted as the percentage difference in the mean level or percentage change per year for a 1 SD increase in PRS.

**^b^** P value is for the difference in the outcome for each 1 SD increase in PRS.

### **Table S10.1** The association between Alzheimer’s disease PRS at *p*≤5x10^-8^ and fasting glucose

| **Outcome** | **Mean difference in glucose (95% CI)** | **Mean difference in glucose (95% CI) per 1 SD higher PRS** | **p ^a^** |
| --- | --- | --- | --- |
| **Glucose** |  |  |  |
| *Age 7yr (mmol/l)* | 4.10 (4.08, 4.13) | 0.01(-0.01, 0.03) | 0.19 |
| Change 7-15yr (mmol/l/yr) | 0.14 (0.13, 0.14) | 0.001 (-0.002, 0.003) | 0.67 |
| Change 15-18yr (mmol/l/yr) | -0.10 (-0.11, -0.09) | -0.01 (-0.02, -0.0001) | 0.05 |

Change/yr, change per year; CI, confidence interval; mmol/l, millimole per litre; mmol/l/year, millimoles per litre per year; SD, standard deviation; yr, year(s).

**^a^** P value is for the difference in the outcome per year where year is indicated for each 1 SD increase in PRS.

Table S10.2 The association between Alzheimer’s disease PRS at *p*≤5x10^-8^, fasting triglycerides and cholesterol

| **Outcome** | **Mean difference in lipids**  **(95% CI)** | **Mean difference in lipids**  **(95% CI) per 1 SD higher PRS** | **p^c^** |
| --- | --- | --- | --- |
| **Triglycerides** |  |  |  |
| *Birth* | -0.66 ln(mmol/l) (-0.69, -0.64)**^a^** | -0.07% (-1.67, 1.54)**^b^** | 0.93 |
| Change 0-9yr | 0.08 ln(mmol/l/yr) (0.08, 0.09) **^a^** | 0.03%/yr (-0.20, 0.26)**^b^** | 0.81 |
| Change 9-18yr | -0.04 ln(mmol/l/yr) (-0.05, -0.04) **^a^** | -0.003%/yr (-0.19, 0.18)**^b^** | 0.97 |
| **Non-HDL-c** |  |  |  |
| *Birth (mmol/l)* | 1.31mmol/l (1.28, 1.34) | 0.005 mmol/l (-0.01, 0.02) | 0.58 |
| Change 0-9yr (mmol/l/yr) | 0.20 mmol/l/yr (0.20, 0.21) | -0.002 mmol/l/yr (-0.01, 0.001) | 0.27 |
| Change 9-18yr (mmol/l/yr) | -0.07 mmol/l/yr (-0.08, -0.07) | 0.002 mmol/l/yr (-0.001, 0.004) | 0.13 |
| **HDL-c** |  |  |  |
| *Birth (mmol/l)* | 0.56 mmol/l (0.54, 0.57) | 0.003 mmol/l (-0.01, 0.01) | 0.48 |
| Change 0-8yr (mmol/l/yr) | 0.12 mmol/l/yr (0.11, 0.12) | -0.0002 mmol/l/yr(-0.002, 0.001) | 0.83 |
| Change 8-18yr (mmol/l/yr) | -0.02 mmol/l/yr (-0.02, -0.02) | -0.0002 mmol/l/yr (-0.001, 0.001) | 0.76 |

Change/yr, change per year; CI, confidence interval; ln, natural log; HDL-c, high-density lipoprotein; mmol/l, millimole per litre; mmol/l, millimole per litre per year; %/yr, percentage per year; PRS, polygenic risk score; SD, standard deviation; yr, year(s).

**^a^** Triglycerides were transformed using the natural log. All values are in log form.

**^b^** The difference in triglycerides per 1 SD higher PRS is back transformed from the log scale for ease of interpretation and is interpreted as the percentage difference in the mean level or percentage change per year for a 1 SD increase in PRS.

**^c^** P value is for the difference in the outcome per year where year is indicated for each 1 SD increase in PRS.

Table S10.3 The association between Alzheimer’s disease PRS at *p*≤5x10^-8^ and fasting CRP

| **Outcome** | **Mean difference in CRP (95% CI)** | **Mean difference in CRP (95% CI) per 1 SD higher PRS** | **p^c^** |
| --- | --- | --- | --- |
| **CRP** |  |  |  |
| *Age 9yr* | -1.25 ln(mg/l) (-1.31, -1.18)**^a^** | -0.01% (-3.73, 3.72)**^b^** | 1.00 |
| Change 9-18yr | 0.10 ln(mg/l/yr) (0.09, 0.11)**^a^** | 0.10%/yr (-0.50, 0.71)**^b^** | 0.74 |

Change/yr, change per year; CI, confidence interval; CRP, c-reactive protein; ln, natural logarithm; %/yr, percentage per year; PRS, polygenic risk score; SD, standard deviation; yr, year(s).

**^a^** CRP was transformed using the natural log. All values are in log form.

**^b^** The difference in CRP per 1 SD higher PRS is back transformed from the log scale for ease of interpretation and is interpreted as the percentage difference in the mean level or percentage change per year for a 1 SD increase in PRS.

**^c^** P value is for the difference in the outcome per year where year is indicated for each 1 SD increase in PRS.
